# Insights from sourdough redefine the domestication landscape of baker’s yeast

**DOI:** 10.64898/2026.04.29.721752

**Authors:** Margot Ruffieux, Nathan Brandt, Alexxis Guttierez, Benjamin E. Wolfe, Robert R. Dunn, Caiti Smukowski Heil

## Abstract

While the domestication of plants and animals is widely recognized for its role in the rise of human civilization, humans have also cultivated microbes over millennia to produce food and beverages. One microbe in particular, *Saccharomyces cerevisiae*, is associated with a wide variety of human-fermentation environments, including wine, beer, and notably bread, such that it is often referred to as “baker’s yeast.” To better illuminate the domestication history of baking associated yeast, we isolated 38 *Saccharomyces cerevisiae* strains from sourdough starters donated by bakers throughout North America and compared them to thousands of *S. cerevisiae* isolates from a variety of wild and human-fermentation environments. We identified 6 major clades with two primary domestication hubs, Mediterranean liquid-state fermentation and Asian solid-state fermentation, diverging across Eurasia that gave rise to human-associated lineages. Population genomic analyses demonstrate that *S. cerevisiae* strains found in sourdough starters are genetically distinct from commercial baking strains and do not come from the surrounding wild environment. Our results show that sourdough yeast strains are closely related to each other and have shared ancestry with strains isolated from various Asian solid state grain fermentations such Japanese sake, Asian rice wines, Chinese distilled spirits (baijiu), and Chinese steamed bread (mantou). We found evidence of significant admixture throughout *S. cerevisiae* populations, including baking-associated lineages, likely facilitated by human activity. Pangenome gene content largely captures *S. cerevisiae* traditional genomic sequence-based population structure and reflects human cultural practices, with differences in gene content and copy number between baking associated strains and other groups. Overall, we show that many generalized hallmarks of domestication, such as genome contraction, loss of genetic diversity, and lack of niche expansion, are not universal features of *S. cerevisiae* domestication, and that baking-associated yeasts have a complex evolutionary history heavily shaped by human culture.

## INTRODUCTION

In addition to the domestication of plants and animals, humans have sustained various fermentations of plant and animal products, facilitating the adaptation and diversification of bacteria, yeasts, and molds to human-associated environments. Evidence of these fermentations dates back over ten thousand years^1–6^, is hypothesized to be far older^7–9^, and has allowed improvements in safety, stability, texture, and flavor of food and beverages^10–15^. In the past decade, the rise of microbial genomics has shed light on how domestication has shaped the genomes of some of these fermentation microbes^16–35^. These studies have demonstrated that microbial domesticates can show reduced genome size, reduced genetic diversity, reduced sexual reproduction, reduced niche breadth, and an increase in copy number alterations and ploidy^22,25,26,29,35–45^. Some of these changes are akin to those seen in plant and animal domesticates^46–49^.

The yeast *Saccharomyces cerevisiae* is one of the most important human-associated microbes, responsible for the production of bread, beer, wine, and other products. Significant population genomics efforts have begun to reveal the biology of temperate forest-associated natural populations^17,21,26,50–62^ and commercial yeast strains used at scale in brewing and winemaking^28,39,63–72^. However, a knowledge gap exists at the interface of these environments, particularly in the evolutionary transitions of yeasts from being wild forest dwellers to their current roles powering wineries, distilleries, breweries, bakeries, and bioethanol production facilities. Commercially available pure-culture yeasts were not available until very recently in human history^73–77^. Instead, sourdough, qu, and variations of “old dough” fermentation starters were some of the sole sources of yeast available for intentionally seeding bread and beer fermentations for millennia^78–84^. Although we do not have yeasts from the earliest versions of fermented doughs, if we consider the traditional starters employed in homes and communities around the world today, we may have a glimpse into thousands of years of yeast and human coevolution.

There are two primary breadmaking practices: adding commercially available baking yeast (*S. cerevisiae*) to dough, or the use of a sourdough starter. Sourdough starter cultures are composed of flour, water, and microorganisms, which serve as the catalyst for dough fermentation^82,85^. Lactic and acetic acid bacteria in starter cultures contribute acidity and flavor^86–88^. Yeasts convert the sugars found in flour into ethanol and carbon dioxide, thus causing dough to rise. Throughout the breadmaking process, the microbial population composition fluctuates in response to selection pressures imposed by the baker and bread dough environment, such as osmotic stress, metabolism of complex sugars, selection for aromas and tastes, and competition with a mixed microbial community^79,85^. Previous studies have suggested commercial and sourdough baking practices apply different selection pressures, and that the yeasts found in those environments belong to separate groups with distinct domestication trajectories^26,44,89^.

Although the origins of sourdough microbes remain poorly understood, spontaneous sourdough starters are often discussed as fermentations formed from wild bacteria and yeasts^90^. A reigning hypothesis for microbial assembly at the community level in sourdough fermentations has been one in which sourdoughs are composed of the microbes that colonize dough in the greatest abundance from flour or other local sources^91–98^. While this is possibly the case for many microbes found in sourdough starters^99^, global sampling efforts of *S. cerevisiae* have clearly found that population structure in *S. cerevisiae* is driven by ecological and functional specialization rather than by geographic origin. *S. cerevisiae* populations are associated with specific environments, including temperate forests, wine, olives, beer, dairy, sake, baijiu, and cacao^20,42^. For instance, a distinct population of *S. cerevisiae* strains found in dairy fermentations carries unique alleles in galactose genes that have allowed those strains to specialize in the milk environment where most other populations could not thrive^100^. Much like dairy strains, many of these populations correlate to specific human activities, and thereby reflect the ongoing role of human cultural practices in *S. cerevisiae* domestication^42,101^.

In an effort to unveil a more comprehensive picture of *S. cerevisiae* domestication, we focus on strains associated with both commercial and sourdough baking practices. We leveraged samples taken by The Global Sourdough Project^88^, a collaborative, community science and society project. On the basis of those samples, we isolated *S. cerevisiae* strains from a subset of sourdough starters donated by bakers and generated high coverage short-read whole-genome sequencing. We combined our sequencing data with additional sequencing of 2,908 *S*. *cerevisiae* isolates^16,17,20,21,26,28,39,42,43,54,59,60,62–64,66–71,102–109^ from a wide range of ecological niches across the globe to conduct large-scale comparative genomics analyses. Drawing on historical, genetic, and ecological records, we uncovered several new populations of dough fermentation yeasts and found evidence for two primary domestication hubs that contributed to further niche expansion and sub-specialization events through admixture. Our results indicate that sourdough yeasts share common ancestors with present-day Asian solid-state fermentation populations and are more closely related to each other than to strains isolated from other fermentation environments. In teasing apart evolutionary relationships among diverse baking yeast populations, this work expands our growing understanding of the intimate, and time-honored relationships between humans and yeasts.

## RESULTS

We selected 50 sourdough starters from over 500 starters donated by bakers through The Global Sourdough Project^88^*. S. cerevisiae* was the most commonly identified yeast species in the Global Sourdough Project; other *Saccharomyces* species were rarely identified. Of the 50 starters we selected, 45 had ITS sequences matching *S. cerevisiae*, 3 matched *S. uvarum*, and 2 contained both *S. cerevisiae* and *S. uvarum*, with these five starters representing all starters containing *S. uvarum*. We successfully isolated 38 *S. cerevisiae* strains, one strain of *S. uvarum*, and one hybrid strain of *S. uvarum* with heterozygous introgression from *S. eubayanus,* from our curated set. Of the *S. cerevisiae* isolates, 34 were obtained from sourdough starters originally started by individuals, as opposed to starters purchased commercially (Figure 1A, Table S1). A majority of the 38 sourdough starters that yielded *S. cerevisiae* were fed unbleached wheat flour, but several were fed rye, whole wheat, and bleached wheat flour (Figure 1B, Table S1). Most of the starters were relatively young (26 =< 5 years old), however, seven starters were reported to be between 12 and 34 years of age, and three were over 100 years old (Figure 1C, Table S1). While isolates predominantly came from sourdough starters located throughout the United States, seven were donated by bakers outside of the US (4 from Canada, 1 from the United Kingdom, 1 from Italy, and 1 from Australia) (Figure 1D, Table S1). In addition to the sourdough samples, two locally available commercial baking yeasts and two commercial brewing yeasts were also sequenced (Table S1).

**Figure 1.**
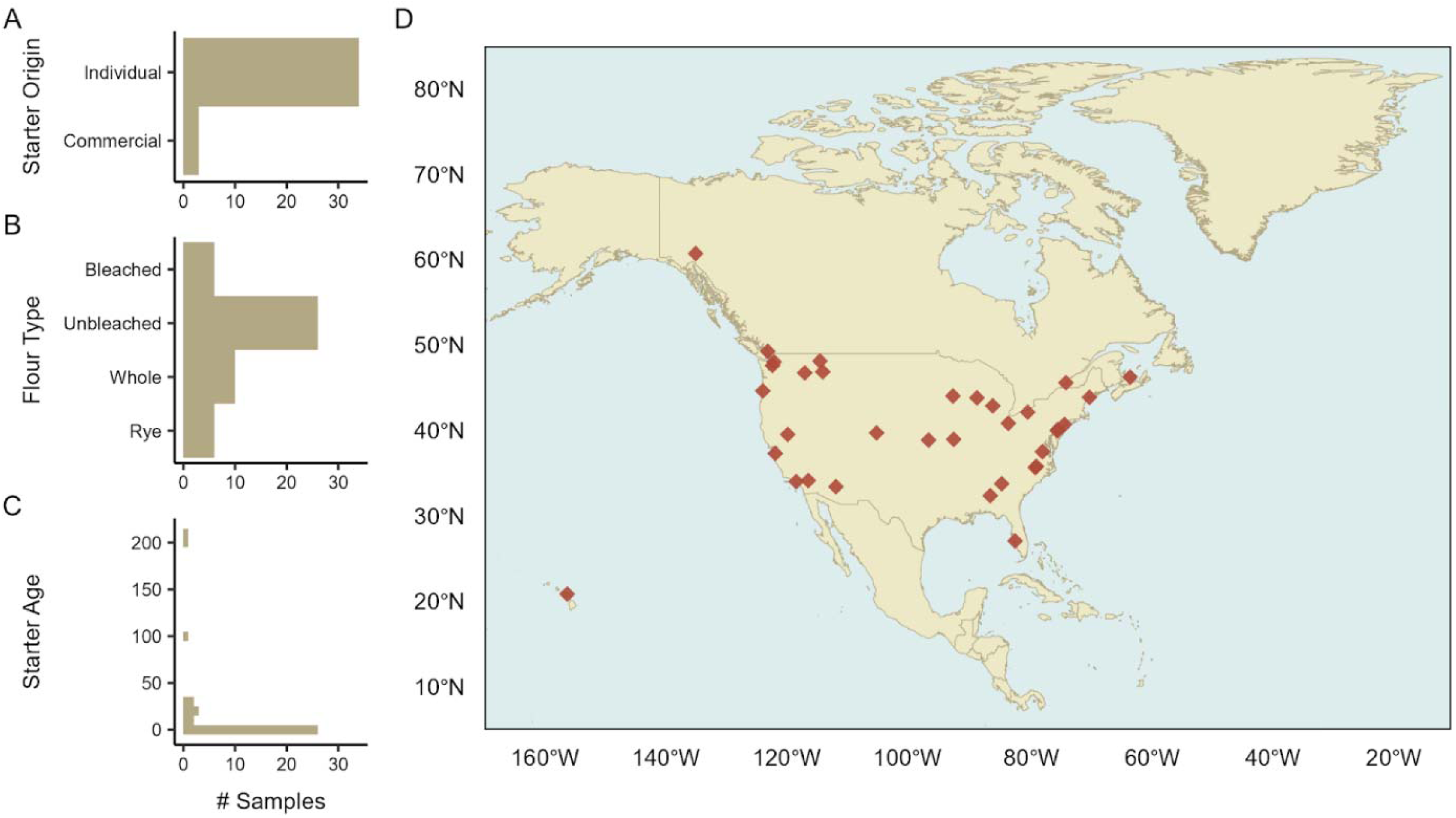
Information about the sourdough starters donated by home-bakers in North America from which *S. cerevisiae* was isolated. Number of samples from sourdough starters A) originally created by individuals versus a starter purchased commercially, B) fed with rye, whole wheat (whole), unbleached wheat (unbleached), and bleached wheat (bleached) flour, C) broken down by reported starter age. D) Map depicting geographic origin of sourdough starters donated by home-bakers in North America from which *S. cerevisiae* yeast were isolated and cultured. Each red diamond represents a sourdough starter. Three starters from the United Kingdom, Italy, and Australia were included in the dataset but are not represented on the map.

To explore the genetic relationships between sourdough yeasts and previously identified *Saccharomyces cerevisiae* populations, the data from the sourdough yeasts were combined with sequencing and variant calls for an additional 2,908 *S. cerevisiae* isolates^16,17,20,21,26,28,39,42,43,54,59,60,62–64,66–71,102–109^ from a wide range of ecological niches (Table S2). The resulting dataset contained genotypes for 2,950 samples and was used to infer a maximum likelihood-like phylogenetic tree from 1,877,031 genome-wide SNPs (Figure 2A, Figure S1). The dataset was further filtered, and population structure was examined for 2,987 samples with FastMixture ancestry prediction (Figure 2B, Figures S2-S3) and principal components analysis (Figure 2C, Figures S4-S5).

**Figure 2.**
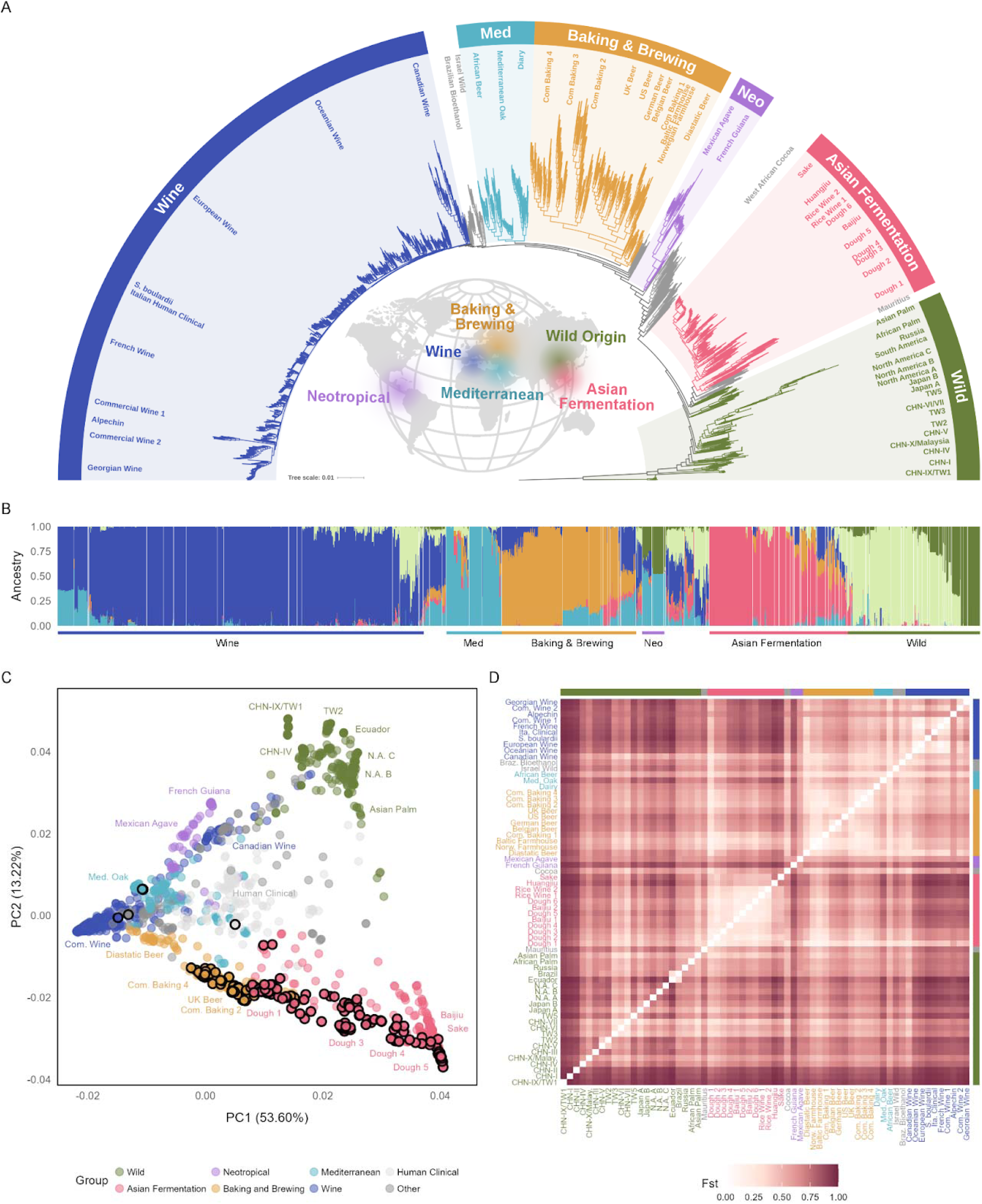
Global phylogeny and population structure of *S. cerevisiae* yeasts. A) Maximum-likelihood-like phylogenetic tree inferred from 1,877,031 genome-wide variant sites in 2950 samples. Branches are colored according to broad groups informed by ecological and evolutionary origins. The world map at the center depicts simplified geographic hubs of putative group origins for major lineages. See Figure S1 for bootstrap support. B) Population structure inferred with FastMixture program for a K value of 6 (Figure S2 and S3). The x-axis is individual samples, grouped by population ordered according to appearance in the phylogeny. The y-axis is the estimated proportion of ancestry for each isolate under the assumption of 6 ancestral populations. C) Scatterplot of the first two principal components (Figure S5 for additional principal components) obtained from principal component analysis run on genome-wide biallelic SNPs. Each point represents an isolate, and isolates are colored according to the grouping scheme and branch colors in phylogeny in panel A. Strains isolated from dough fermentation environments are outlined in black. Several populations depicted in the phylogeny are annotated with text labels in the scatterplot for clarity. D) Heatmap of pairwise *F_ST_* for all pairs of populations identified through phylogenetic analysis (Table S3). Populations are labeled on both the x and y axes, and population labels are colored by the broad groups depicted in panels A-C. *F_ST_* values are represented by a continuous color scale, with low values in light shades and high values in dark shades.

### Two yeast domestication hubs are associated with an east-west divergence across Eurasia

Phylogenomic and population structure results recovered many of the same populations found in prior studies and support previous observations that human-associated *S. cerevisiae* populations are distinct from wild populations and reflect human cultural practices^16,19,20,101,108^ (Figure 2A-C). All three population structure analyses revealed six superclades, 1) Wild, 2) Asian Fermentation, 3) Neotropical Fermentation 4) Baking and Brewing, 5) Mediterranean Fermentation, and 6) Wine Fermentation (Figure 2A-C, Figures S2-3). Five of the six major groups are associated with human fermentation environments. The human-associated groups appear to arise from two primary domestication hubs associated with an east-west divergence across Eurasia. Admixture between these two groups, or between these groups and wild lineages, forms additional subpopulations (Figure 2A-D). The separation between the two main domestication hubs explains 53.6% of genotype variation (PC1, Figure 2C) and roughly corresponds to a Western Eurasia branch composed of Mediterranean/Wine fermentation lineages and an Eastern Eurasia branch composed of Asian Fermentation/grain fermentation lineages. In line with prior observations, the two domestication hubs are associated with fermentation substrates that differ in viscosity and water content: the Eastern group with solid-state grain ferments and the Western group with liquid-state fruit ferments^106^. Strains comprising the Western group are found in wine, cider, olive brine, dairy, and traditional African beers, and the majority of strains are isolated from Europe and the Mediterranean Basin, as well as vineyards and wine fermentations in Oceania and North America where viticulture has spread more recently^64,68–70^. The Eastern group is geographically centered in Asia, specifically what are now China, Japan, and Southeast Asia, and consists of strains isolated from grain fermentations that make up the previously established Asian Fermentation superclade^20,26,106,108^ (Figure 2A). Isolation substrates for the Eastern group include rice ferments (sake, huangjiu, rice wine), sorghum ferments (baijiu), and wheat ferments (dough, sourdough starters, mantou, qu). Across the entire phylogeny, population pairwise *F_ST_* estimates were high, ranging from 0.05027 to 0.97192 (Figure 2D). This is consistent with previous findings that over 90% of species-wide SNPs are present at low allele frequencies (MAF<0.1)^20^ and is a pattern indicative of population expansion following a bottleneck, a common scenario during domestication^110^.

### Sourdough yeasts form populations of dough fermentation strains and are closely related to populations associated with other grain fermentations

The second PCA (PC2, Figure 2C) axis further differentiates strains associated with grain fermentations from strains found in vineyards and forests, with an overarching pattern of separating wild populations from human-associated ones. A large proportion of sourdough and dough fermentation isolates from North America and Europe are closely related to Asian Fermentation lineages and form populations on the Asian Fermentation phylogenetic branch^26,44^ (Figure 2A). Phylogenomic analyses revealed six populations (named Dough 1-6) associated with dough fermentation environments, several of which encapsulated and expanded on previously identified mantou (dough fermentation) populations^26^ and further resolved an ambiguous region of the *S. cerevisiae* phylogeny coined “Mosaic Region 3”^20^ (Figure 2A). Dough 1 and Dough 2 populations are both composed of an amalgamation of dough fermentation strains, which include a combination of previously identified mantou populations alongside newly added sourdough genomes with geographic origins spanning Asia, Europe, and North America^26,44^ (Figure 2A, Table S2). Dough 3, Dough 4, and Dough 5 are all composed of Chinese mantou isolates collected from homes throughout China^26^. The final dough fermentation group identified, Dough 6, is closely related to Asian rice wine populations (Rice Wine, Huangjiu, Sake), and is characterized by samples isolated from sourdough starters donated by bakers throughout North America. North American sourdough isolates make up 16 out of the 23 strains in the Dough 6 population (Figure 2A, Table S2).

Two dough fermentation populations, Dough 1 and Dough 2, appear to be early branching members of the Asian Fermentation superclade. However, it is possible that their “basal” phylogenetic positioning and long branch lengths are an artifact of heterozygosity and admixture, as has been suggested for the European Farmhouse yeasts^71^ (Figure 2A, 2B). Previous work has shown that many human-associated lineages are admixed. Our phylogenomic and population structure results indicate that dough groups exhibit varying proportions of ancestry from other superclades (Figure 2) and have low pairwise *F_ST_* values with populations outside of the Asian Fermentation group (Figure 2D, Table S3). Overall, these results support admixture and gene flow among divergent *S. cerevisiae* lineages as evolutionary forces that shaped the genomes of sourdough strains, particularly those in Dough 1 and Dough 2 populations.

### Commercial baking yeast are genetically distinct from sourdough populations and share a common origin with European beer yeasts

Four commercial baking populations (Commercial Baking 1-4) were identified within the Baking and Brewing superclade (Figure 2A). These populations are primarily composed of strains identified as commercially available baking yeasts and bakery samples from across the world. Fifteen strains isolated from sourdough fell within the Baking and Brewing superclade in both the phylogenetic tree and the PCA (Figure 2A, Figure 2C); those samples were tightly clustered with known commercial baking strains and are likely commercial baking yeasts that were either intentionally or unintentionally introduced to the sourdough starter environment. *F_ST_* between Asian Fermentation dough groups and Commercial Baking groups (0.113-0.804, Figure 2D, Table S3), phylogenetic placement, and population structure all support distinct evolutionary trajectories of sourdough and commercial baking yeasts, building upon previous findings from Bigey et. al^44^. Commercial baking isolates are most closely related to brewing isolates from the European beer subclades (Figure 2A, Figure 2C). Shared ancestry inferred through FastMixture analysis, as well as low population pairwise F_ST_ among Baking and Brewing populations (0.050-0.470, Figure 2D, Table S3) and increased ploidy of both commercial baking and brewing strains (Figure 4), suggest that the commercial baking and brewing populations likely share a common ancestor.

**Figure 3.**
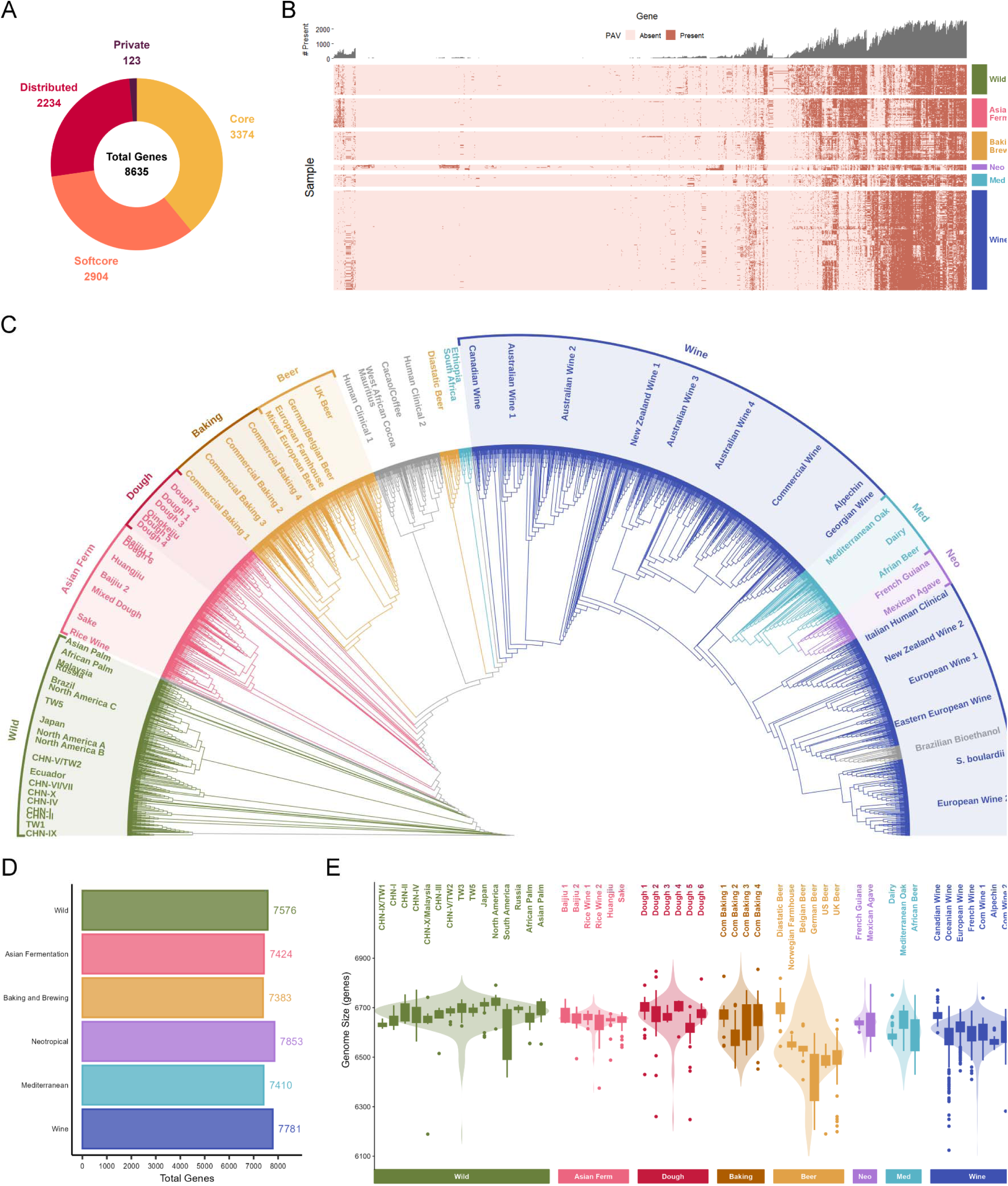
Characterization of pangenomic gene content for 2,913 *Saccharomyces cerevisiae* strains. A) Ring plot of pangenome composition. B) Heatmap of pangenome gene presence/absence variation (PAV) in distributed genes for all isolates. Samples are on the y-axis and are split by broad evolutionary groups. Genes are hierarchically clustered on the x-axis. The bar chart at the top of the heatmap along the x-axis displays the total number of samples (out of 2,913) in which each pangenome gene was detected. C) Phylogenetic tree inferred under a binary model from pangenome gene presence/absence profiles for 2,913 strains. Branches and population labels are colored according to corresponding evolutionary groups identified in Figure 2. See Figure S9 for bootstrap support. D) Bar chart of total pangenome gene diversity for each evolutionary group. E) Violin plot depicting number of genes per genome across groups, overlaid with boxplot highlighting genes per genome broken down by population. Groups are colored according to the broad groups shown in the nucleotide phylogeny and PCA plots of Figure 2, but Dough Fermentation is plotted independently from Asian Fermentation and Baking and Brewing are separated into two groups, Baking and Beer. Significance is reported in Table S9.

**Figure 4.**
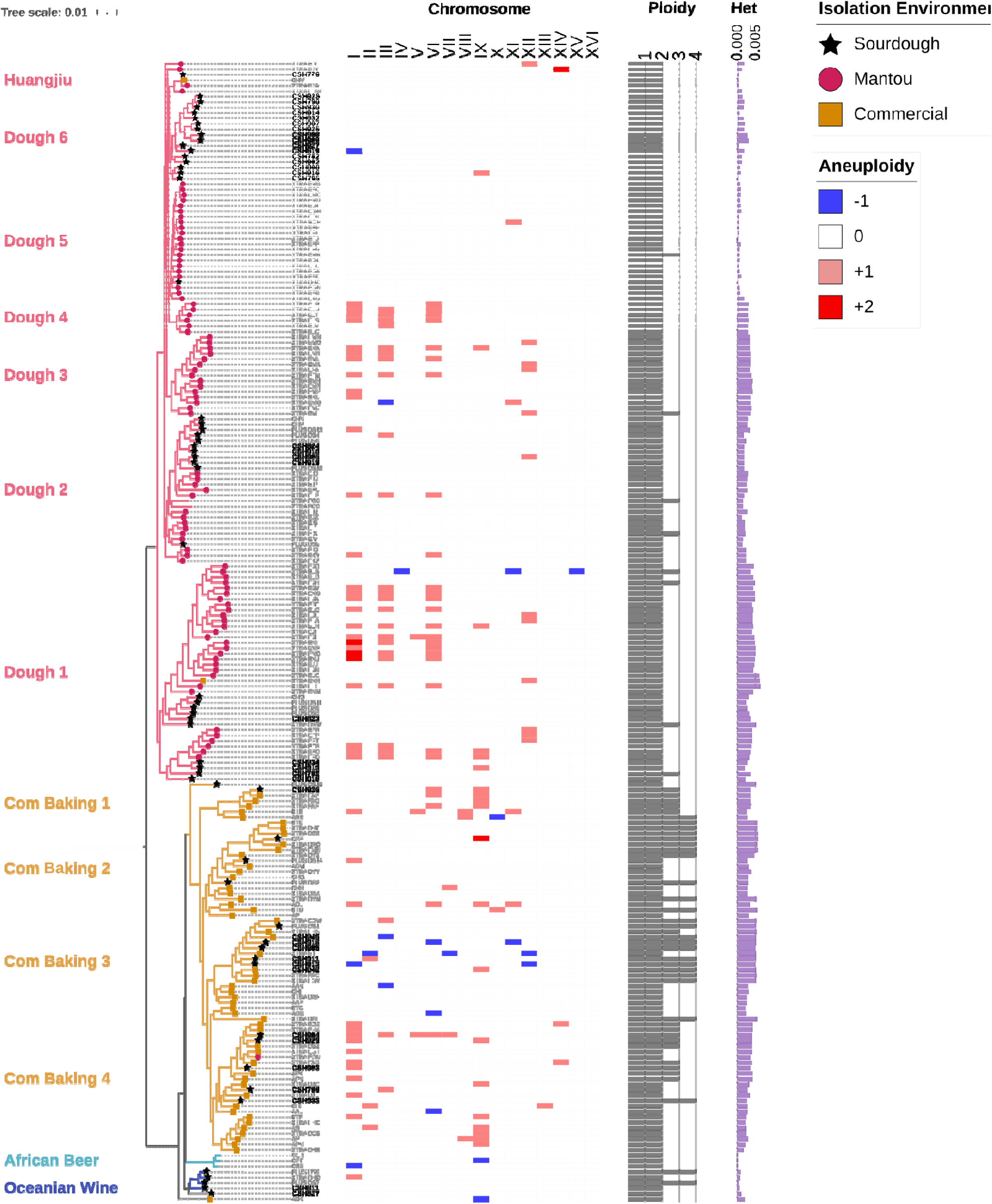
Pruned phylogeny of 209 baking-associated *S. cerevisiae* isolates annotated with isolation environment, ploidy, aneuploidies, and proportion of genome-wide heterozygosity. From left to right, the first annotation layer is labeled by phylogenetic population and colored according to the broad evolutionary group (as per Figure 2). Branch colors also correspond to broad evolutionary groups. Branch symbols indicate the type of baking environment from which each sample was isolated (pink circle: mantou; black star: sourdough; gold square: commercial). Bolded black tip labels signify isolates newly sequenced in this work. Note that samples CHS942 and CSH945 were isolated from commercially available baking yeast. The heatmap depicts chromosomal aneuploidies across 16 nuclear chromosomes, with blue representing chromosomal loss and red chromosomal gain. The black bar chart shows estimated genomic ploidy, and the purple bar chart is genome-wide heterozygosity.

### Pangenome gene content recapitulates evolutionary history

To identify how domestication has influenced gene content, we mapped short reads for 2,913 strains (Table S2) to the gene-based pangenome reference created by Loegler et al.^111^ and called gene copy number and presence/absence variation (PAV). The pangenome reference contained 8,541 genes^111^, of which 8,535 were found to be present in our dataset (Data S1, Data S2). There were 3,374 genes present in every strain, forming the strict core genes (Figure 3A). An additional 2,904 softcore genes were found in over 95% of samples, extending the core and softcore (95%+) to 6,278 genes. There were 2,234 genes identified as distributed genes (>=2 samples, <95% of samples), and 123 genes were categorized as private genes (singletons, present in only one sample) (Figure 3A, Figure S6, Tables S6-S7). Pangenome gene accumulation curves for both core and distributed genes identified the pangenome as a closed pangenome (Figure S6).

We next explored whether pangenome presence/absence reflects traditional evolutionary relationships obtained through nucleotide sequence variation. To do so, we created a heatmap of the 2,234 distributed pangenome genes across samples (Figure 3B). The heatmap of pangenome gene PAVs displayed identifiable patterns among Wild, Asian Fermentation, Commercial Baking & Brewing, Neotropical, Mediterranean, and Wine groups, demonstrating that pangenome gene content variation does reflect broad evolutionary groups (Figure 3B). To determine if pangenome gene PAVs also reflect finer-scale population structure, we inferred a phylogenetic tree using pangenome gene presence/absence under a binary model (Figure 3C, Figure S8). The resulting phylogeny largely recovered the populations, evolutionary groups, and branching patterns observed in the sequence-based phylogeny (Figure 2A), indicating that population structure at both fine and broad scales is captured by pangenome gene content variation (Figure 3C, Figure S9). We also examined the frequencies of pangenome genes across populations and several genes emerged (YDR036C, YX01845, YX00831, and YLR460C, Figure S8) that may be implicated in the separation between east Eurasian and west Eurasian domestication trajectories identified in SNP-based population structure analysis. Much like the population structure recovered by reference-based SNPs, population structure identified by pangenome PAVs separated wild populations from human-associated populations and grouped isolates by fermentation process and environment (Figure 2A, Figure 3B-C). The concordance of population structure reflecting domestication history across SNP and pangenome gene PAV genetic variation demonstrates that evolutionary history is captured at both mutational and gene-content levels, and further underscores that *S. cerevisiae* evolution has been shaped by human culture.

### Genome decay is not consistently associated with domesticated populations

Pangenome gene content analyses in soybeans, common beans, tomatoes, and goats identified gene loss in domesticated populations relative to wild populations, linking genome reduction to domestication in both plant and animal domesticates^47,112–115^. Previous studies on beer yeast and other microbes have also demonstrated that genome decay is associated with microbial domestication^29,39,40^. The so-called “genome reduction hypothesis” of domestication derives from studies of the evolution of vertically inherited symbionts relative to their hosts^116–118^. This loss is expected on the basis of the loss of need for many functions, such as dispersal and metabolism of nutrients reliably provided by the host or ferment. Additionally, repeated selection, genetic bottlenecks, and loss of niche breadth during adaptation to food environments can lead to a reduction in genetic diversity of domesticated microbes^40^. To look for evidence of reduced gene diversity and genome decay as consequences of domestication broadly across *S. cerevisiae* populations, we first calculated total pangenome gene diversity for each of the six major evolutionary groups (Figure 3D, Figure S7). Pangenome gene diversity results differed from expectations based on genome-wide nucleotide diversity, where the wild group had the highest nucleotide diversity (Table S4). Surprisingly, both Neotropical Fermentation and Wine groups harbored greater pangenome gene diversity than all the wild populations combined (Figure 3D, Figure S7).

We then counted the number of pangenome genes present in each strain to serve as a proxy for genome size and visualized the gene counts per-strain by population (Figure 3E, Table S8). Surprisingly, only 35% of pairwise comparisons between wild and human-associated populations yielded a significant difference in genome size, contradicting expectations of genome decay (Dunn Test, p<0.05, Table S9). While we did indeed see that European beer yeast and several Wine yeast populations tended to have fewer genes than their wild counterparts, Baijiu, Dough Fermentation, and some Commercial Baking populations have similar gene counts to wild populations (Figure 3E, Tables S8-S9). Domesticated populations with significant gene loss were generally members of the west Eurasian/liquid-state fermentation group, whereas those with wild-like gene content mostly fell along the east Eurasian/solid-state fermentation domestication trajectory (Figure 2). Although it is possible that some food-associated strains are more domesticated than others (i.e. more isolated or under stronger selection for domestication traits), the Asian Fermentation group also contained highly isolated and industrialized strains (huangjiu/ bioethanol, sake). This discrepancy in genome size mirroring the east-west Eurasian divergence in population structure suggests that genome reduction could be the result of a particular evolutionary path to domestication, rather than a global hallmark of domestication.

### Baking-associated isolates form populations in Asian Fermentation and Baking and Brewing superclades, and have lineage-specific genomic features

To characterize genomic similarities among just the baking strains, a sub-phylogeny of 209 baking-associated strains^20,26,44,67^ was pruned from the global tree and used to visualize shared patterns of isolation environment, aneuploidies, ploidy and heterozygosity, in dough fermentation populations (Figure 4, Table S10). Baking-associated strains were determined by available metadata, where any sample with an isolation source related to dough fermentation, sourdough, bread, baking, bakery, or commercial baking/active dry yeast was included (Table S2, Table S10). The final baking-associated dataset consisted of 90 samples from mantou, 59 samples from sourdough, and 55 samples from commercial baking products (Figure 4, Table S10). For newly isolated and sequenced strains, ploidy was estimated from DNA content by flow cytometry, and aneuploidies were called computationally from BAM files using sequencing read depth (Figure S11).

Baking strains from across the globe form thirteen populations, largely falling in the Asian Fermentation and Baking and Brewing superclades, with specific patterns of ploidy, aneuploidies, and heterozygosity (Figure 4). Sourdough isolates sequenced in this study were predominately placed among the six new dough populations in the Asian Fermentation superclade, notably in the North American Sourdough population (Dough 6). Interestingly, the Dough 1 and Dough 2 populations were each composed of strains isolated from diverse dough ferments spanning three different continents (Table S2). Our results suggest relatively strong “conservatism” of sourdough *S. cerevisiae* populations, wherein sourdough strains tend to derive from closely related dough-associated populations, with some exceptions. For example, two sourdough isolates, one from Australia and another from the United States, were among admixed Australian wine isolates in the Oceanian Wine population. We hypothesize that sourdoughs in those cases resulted from colonization of wine yeasts into new starters. Additionally, bakery isolates from Ghana cluster with other Ghanaian fermentation strains rather than grouping with global dough fermentation or bakery samples. In other words, sourdoughs can form with strains derived from other sources, but these cases are exceptions.

Baking strains in the Baking and Brewing superclade had slightly higher average genome-wide genetic diversity (0.0036) than sourdough strains in the Asian Fermentation superclade (0.0026), possibly reflecting signatures of increased ploidy and admixture (Figure 4, Table S10, Table S4). Commercial baking populations had the most instances of elevated ploidy (13 triploid, 20 tetraploid), a relatively unique feature shared by the strains in the Commercial Baking & Brewing superclade (Figure 4, Table S10). Relatedly, the commercial baking strains had the highest frequency of both chromosomal gain and loss aneuploidies throughout the nuclear genome (Figure 4, Table S6), consistent with previous findings^45^. In contrast, sourdough-specific clades were mostly diploid, and their aneuploidies were generally restricted to chromosome gains on Chromosomes I, III, VI, and XII (Figure 4, Table S10).

### Maltose metabolism and sporulation phenotypes of breadmaking yeasts

Using a subset of representative strains alongside sourdough and commercial baking isolates, we measured growth rate in maltose media as a proxy for the ability to metabolize maltose, a primary complex sugar available in dough fermentation environments (Figure 5, Table S11). A sub-phylogeny was pruned from the full phylogeny in Figure 2 for the phenotyped strains, and both the maltose growth rate data (Table S11) and copy number data for pangenome genes annotated with functions associated with maltose metabolism (Data S3) were plotted along the pruned tree (Figure 5).

**Figure 5.**
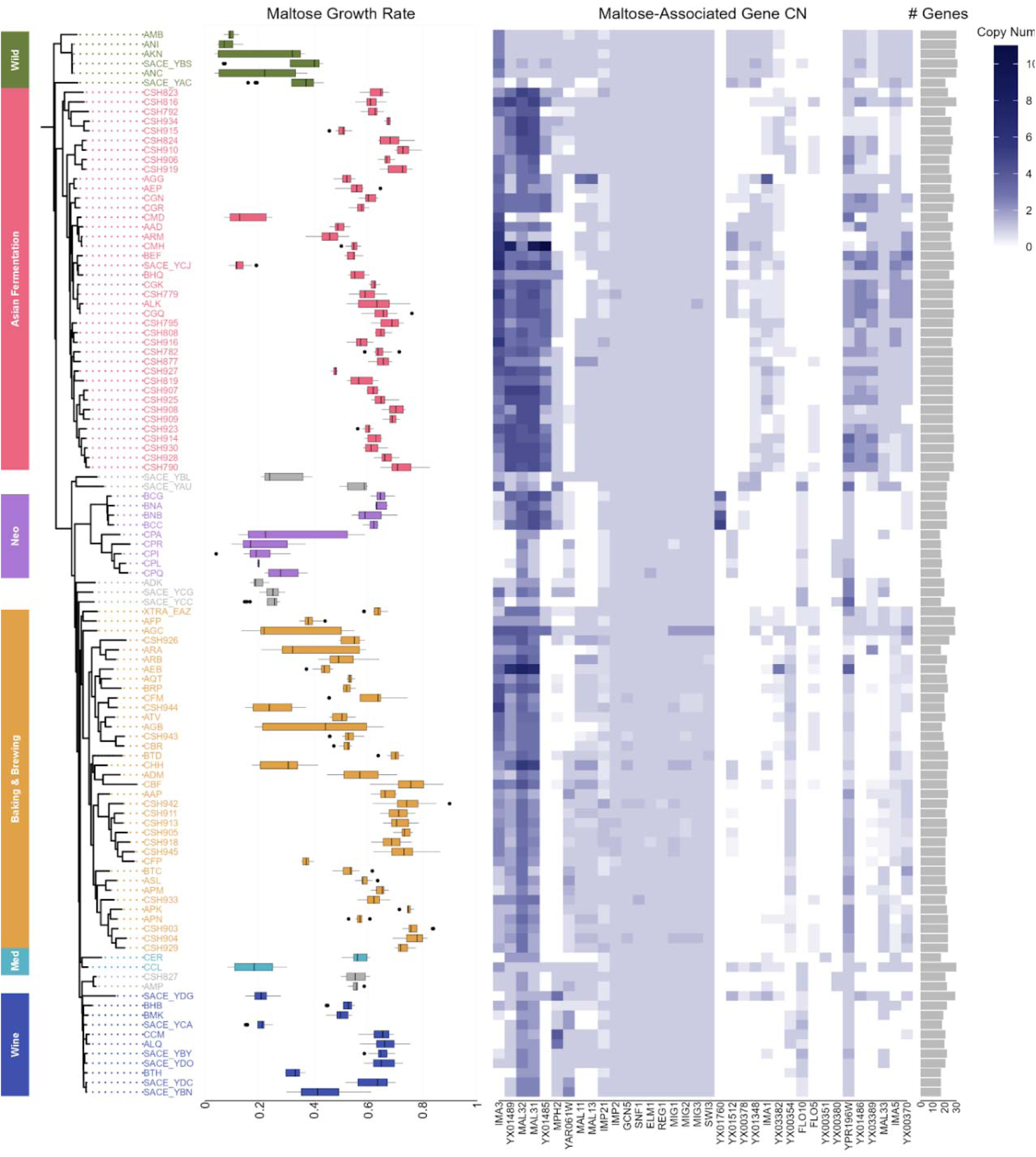
Boxplot of growth rate (r) in maltose media measured for 111 isolates from Wild, Asian Fermentation, Neotropical, Baking and Brewing, Mediterranean, and Wine groups alongside a heatmap of copy number (CN) variation of maltose-associated pangenome genes. Plots are aligned to a phylogeny pruned from the full tree in Figure 2A, with samples on the y-axis. Tip labels and box plot are colored according to broad groups, as previously delineated (Figure 2). The bar plot on the right indicates the number of maltose-associated genes detected for each isolate. The boxplot x-axis is growth rate, the heatmap x-axis is maltose-associated pangenome genes, with their order determined by hierarchical clustering, and the bar plot x-axis is the number of genes present. In the copy number heatmap, gene absence is shown in white, and an increasingly higher copy number is represented monochromatically by a continuous scale of deepening blue.

Although differences in maltose metabolism can be seen across the tree, the total number of maltose-associated genes present in each strain’s genome (rightmost column of Figure 5) was not correlated to maltose growth rate (R^2^=0.009, p=0.324). Wild strains had as many maltose-associated genes as strains found in maltose-heavy environments but did not grow as well in maltose media as Asian Fermentation, Commercial Baking, and Wine strains. Patterns of copy number variation can be seen by clade and maltose growth rate (Figure 5). To determine if copy number of any maltose-associated genes was correlated with maltose growth rate, we calculated phylogenetically corrected spearman correlations using phylogenetic independent contrasts. Copy number of one gene, YX00354, was found to have a significant positive correlation with growth rate in maltose (Figure S12, spearman correlation, p=0.047). YX00354 is a novel gene identified in the pangenome produced by Loegler et al.^111^ that was annotated as a maltose fermentation regulatory protein highly similar to MAL33. Interestingly, the S288C reference version of *S. cerevisiae* MAL33 was not significantly correlated to maltose growth rate (Figure S12), but allelic variation of the MAL33 gene in *S. eubayanus* was recently identified as a driver in strain-level variation of maltose metabolism^119^. These results point to the significant role pangenome gene copy number plays in maltose metabolism^26,44^ and highlight the importance of considering pangenomic variation when connecting genotype and phenotype^111^ (Figure 5, Table S11, Data S3).

In addition to maltose metabolism, a subset of sourdough and commercial baking strains were tested for spore viability and sporulation efficiency, as an inability to complete a sexual cycle is associated with yeast domestication^22^ (Figure S13, Table S12). Sporulation efficiency was highly variable, with some Commercial Baking strains exhibiting near complete sporulation whereas other strains were unable to sporulate. Asian Fermentation sourdough isolates showed intermediate sporulation efficiencies, though overall there was no significant difference in means between the two baking groups in sporulation (two sample t-test, p=0.058). Spore viability was significantly lower in commercial baking strains compared to sourdough strains (two sample t-test, p=0.021), and surprisingly, only tetraploid commercial strains produced viable spores; other ploidies showed no pattern in either baking group.

## DISCUSSION

Here, by expanding *S. cerevisiae* sampling with isolates from North American sourdough starters and building on recent genome projects, we leveraged a large-scale comparative genomics approach to reveal patterns in *S. cerevisiae* evolutionary history and domestication. Through the exploration of wild and human-associated populations spanning continents and cultures of one of the oldest microbial domesticates, this work advances our understanding of the unique partnership between yeast and humans and the genomic hallmarks of domestication at large.

In sampling home-baker sourdough starters from North America, we found new global populations of sourdough yeast, deemed Dough 1-6, and identified their genetic proximity to strains from other Asian solid-state grain fermentations^26,44^. Often, the microbial assembly of spontaneous sourdough starters is discussed as though each instance of a starter independently draws from a diversity of potential yeasts from the substrate and the surrounding environment^97^. Our results revealed that *S. cerevisiae* sourdough yeast do not come from the surrounding wild environment, but rather, from populations of yeast that specialize in dough and grain fermentations. Geographically distant yeast strains from spontaneous dough fermentations are more closely related to other dough strains than they are to geographically proximal strains inhabiting different environments, as expected if sourdough yeasts tend to derive from ecotypes adapted to this environment. This is demonstrated by the North American Sourdough population (Dough 6), which is much more closely related to other dough and grain fermentation strains isolated throughout Asia than strains isolated from oak trees, beer, or wine in North America. Together, this suggests that *S. cerevisiae* sourdough yeasts are passed from ferment to ferment, but it remains unclear how closely related strains become so geographically dispersed. There is currently no strong evidence to support that *S. cerevisiae* in sourdoughs comes from cereal grains or flour, and instead house microbiota may be the key source in bakeries^94,98^. More work is needed to investigate strain level diversity in bakeries and homes to determine potential sources, though humans are clearly important in dispersal.

Our phylogenomic and population structure results identified six major evolutionary groups, of which five out of six are composed of strains and populations primarily associated with humans. An overarching pattern distinguishing the various human-associated groups was fermentation process, as demonstrated by the grouping structure of Asian Fermentation, Baking and Brewing, and Wine reflecting solid-state grain fermentation, liquid-state grain fermentation, and liquid-state fruit fermentation respectively. We found evidence for two domestication centers that generally follow an east-west divergence across Eurasia and differ by substrate hydration levels^106^, with finer structure of domesticated yeast clades reflecting human cultures. In the east Eurasian branch, Sake, Rice Wine, Baijiu, and Dough Fermentation populations from the Asian fermentation group revealed populations specific to regional variations in cultures and traditions. In the west Eurasian branch, African Beer, Dairy, Alpechin, and Wine clades reflect yeast population diversification in tandem with the spread and differentiation of human fermentation cultures throughout the Mediterranean Basin and Europe. More recently, advancements in technology and globalization have steered the evolutionary course of *S. cerevisiae* commercial baking yeast through the development of Commercial Baking populations that branched off from European beer populations. We corroborate previous findings that there are two different evolutionary trajectories for yeast used in baking, with sourdough strains placed in the Asian Fermentation superclade and industrial baking yeast placed in Baking and Brewing superclade^26,44^. Our results support a common origin between Commercial Baking yeast and European ale yeasts^19,42,67^. Together, Commercial Baking yeasts and European Beer yeasts form the Baking and Brewing phylogenetic superclade^19^, and isolates within the Baking and Brewing superclade share genomic characteristics that are relatively unique to that group. Although polyploidy has been implicated as a broad symptom of domestication in *S. cerevisiae* yeasts, the Baking and Brewing yeasts are typically tetraploid, while very few isolates outside of that group are^20,22,28,39,108,120^. Phenotypic analysis of spore viability also identified a distinct pattern in commercial baking strains from the Baking and Brewing group that differed from Asian Fermentation strains. Of the strains tested for spore viability, only tetraploid commercial baking strains produced viable spores. The Baking and Brewing yeasts also have uniquely high heterozygosity, share similar admixture profiles, and the pairwise *F_ST_* among Baking and Brewing populations reveals low differentiation within the superclade.

Much like human populations, yeasts have a complex evolutionary history shaped by past and ongoing gene flow. Our results support previous findings of repeated role of admixture in *S. cerevisiae* evolution, including an admixed origin of Commercial Baking and Brewing yeasts^67^. Similarly, sourdough yeasts exhibit genomic signatures of admixture. Evidence from ancestry profiles, phylogenetic and PCA placement, and low *F_ST_* values with populations in non-baking groups denote the presence of gene flow or admixture events in Dough 1 and Dough 2 populations. For that reason, the precise origins of sourdough strains remain unclear. It is possible that the spontaneous fermentation dough environment facilitates admixture, much like has been previously suggested in spontaneous wine fermentations^68–70^. Another possibility is that sourdough yeasts arose from a historical admixture event, and through dispersal, isolation, selection, and drift, various subpopulations have retained differing proportions of ancestral alleles. In addition, sourdough yeasts may have experienced more recent gene flow with non-sourdough populations or yet undiscovered populations. None of these hypotheses are mutually exclusive, and it is likely that some combination of these forces is at play. Environmental and archeological sampling in unsampled regions with historical connections to wheat domestication and bread cultures, like West Asia, may help untangle these past and present evolutionary dynamics. More generally, we see a trend that human-associated populations experience increased gene flow compared to wild populations, supporting prior observations^121^. Although yeasts are closely associated with and dispersed by humans^62,64,69,122^, *S. cerevisiae* populations have much higher pairwise *F_ST_* (up to 0.972) than human populations (less than 0.3)^123^. Instead, *F_ST_* among *S. cerevisiae* populations is more akin to that of other wild *Saccharomyces*^124,125^ species despite *S. cerevisiae* evolutionary history having been greatly altered by domestication.

Zooming out, our results have broad implications for our understanding of *S. cerevisiae* evolution and domestication syndrome. Interrogation of pangenomic gene content revealed that pangenome gene presence/absence variation does largely recapture *S. cerevisiae* population structure identified through traditional reference-based SNP variant approaches^19,20,108^. Much like sequence-based population structure, pangenome PAV-based population structure was primarily associated with fermentation process and environment, including clear differences between sourdough and commercial baking strains. Our analysis of pangenome gene copy number for maltose-associated genes suggests that pangenome gene copy number, in addition to presence/absence, plays a role in adaptation to baking environments. Copy number variation and gene family expansion of maltose associated genes are now recognized as a key signature of *S. cerevisiae* domestication to wheat and barley based solid-state and liquid-state fermentations^26,28,42,44,126^. Distributed genes have been previously shown to contribute to transcriptional diversity and fermentation-specific patterns of expression^128,130^, so investigation into particular PAV patterns and newly identified gene functions specific to fermentation types is an exciting area for future study.

Despite the prominent role domesticated microbes play in human society, microbial domestication is understudied and remains poorly understood. Previous work has outlined several features associated with microbial domestication, including distinct subpopulations, gene loss, genome decay, increased ploidy and aneuploidy, increased copy number, loss of sexual reproduction, specialized metabolism, and lack of niche expansion^26,29,38–41,44,106^. Here, we show that while some of these hallmarks of domestication hold true for particular lineages, many *S. cerevisiae* domesticated populations have not experienced a decrease in gene content compared to wild populations. Similarly, most human-associated populations have higher heterozygosity and equally high or higher genome-wide nucleotide diversity as wild populations. Human-associated populations have also experienced increased gene flow as a consequence of occupying human fermentation environments, though relaxed selectionmay be leading to loss of sexual reproduction. While in *S. cerevisiae* we do see a clear and consistent pattern of distinct subpopulations specific to ferment type, the formation of those subpopulations is primarily associated with niche expansion as a result of human association. Our results challenge the assumptions that genome decay, reduced outcrossing, decreased genetic diversity, and lack of niche expansion are general hallmarks of microbial domestication. Perhaps traditional fermentation practices and spontaneous ferments are key to maintaining and promoting genetic diversity in domesticated yeasts by facilitating encounters of strains from disparate populations. Overall, our results shed light on the evolution and domestication of yeasts in traditional and commercial human fermentations, and in doing so, unveil previously hidden dimensions of human and microbial partnerships.

## METHOD DETAILS

### Isolation & Identification of S*. cerevisiae* from Sourdough Starters

A collection of 50 sourdough starters was obtained from a previously published study of The Global Sourdough Project ^88^. Sourdough starters contained a mixed microbial community, so starters were prioritized to 1) contain a large proportion of ITS locus amplicon reads matching *S. cerevisiae*, 2) maximize *S. cerevisiae* haplotype diversity of the ITS locus, 3) sample from a broad geographic distribution in North America, and 4) obtain any starters that contained ITS locus amplicons matching to *S. uvarum*. To identify *S. cerevisiae*, individual colonies were screened using colony PCR with primers CSH389 (CGGGTAACGAAGAATCGACTCTCGC) and CSH390 (GCGGAATAACCTAACACGTGGAGG). These primers were used in conjunction with yCSH453 (*Lachancea thermotolerans*) as a negative control. Five colonies per sourdough starter were initially screened. If no *S. cerevisiae* was identified, additional colonies were screened. In total, we were able to successfully isolate 38 *S. cerevisiae* strains from 47 sourdough starters (Table S1). Failure to recover *S. cerevisiae* from a subset of the starters may be due to discrepancies between amplicon read abundance and actual culture abundance, or issues with primer specificity. We additionally isolated 1 *S. uvarum* strain and 1 *S. eubayanus x S. uvarum*hybrid with primers 39_uva_chrXIII_194496F (GATTCACCACGCCTAAGCTCTTGAG) and 39_uva_chrXIII_194496R (GTTGTCAAGGAATTGCCAAAGCC) from 5 sourdoughs that had an ITS match to *S. uvarum* (labeled as *S. bayanus*). These 5 sourdoughs were the only sourdoughs in the collection^88^ that had a *Saccharomyces* species besides *S. cerev*is*iae* according to the ITS locus. Isolated strains were grown in YPD media + ampicillin (100 µg/mL) + kanamycin (100 µg/mL) and archived in 15% glycerol at -80°C.

### DNA Extraction, Library Construction, and Whole-Genome Sequencing

For all samples, genomic DNA was extracted using a modified Hoffman-Winston protocol^127^. SYBR Green I was used to measure genomic DNA concentration, and the input for each sample was standardized to 250 ng. Libraries were constructed using the Illumina DNA Prep Kit. The concentration of purified libraries was then measured again using SYBR Green I, and libraries were pooled to balance their concentrations. The pooled library was sequenced with paired-end reads (2 x 150 bp) on an Illumina NovaSeq (Duke University Sequencing and Genomic Technologies).

### Read Mapping and Variant Calling

Illumina short-reads for 40 newly sequenced isolates (Table S1), along with 1,333 publicly available shortreads^20,26,44,60,62,67,104,107^ (Table S2) were aligned to the *S. cerevisiae* S288C R64-3-1 reference genome^129^ (downloaded from the Saccharomyces Genome Database^131^) with BWA-mem version 0.7.17^132^ using default parameters. Alignment files were quality-checked with FastQC version 0.11.9^133^ and then sorted and indexed using Samtools version 1.13^134^. Picard version 2.25.6^135^ was used to mark and remove duplicate reads and assign read groups. Samtools was then used to filter alignments, excluding reads with a mapping quality below 20, unmapped reads, secondary alignments, and reads failing quality control. The filtered BAM files were again sorted and indexed with Samtools to produce the final alignment files used for downstream analysis. Variants were called with the HaplotypeCaller function in the Genome Analysis Toolkit (GATK) version 4.5.0^136^, producing all-sites GVCFs for each sample. The resulting GVCF files, along with previously published GVCF files for an additional 1,610 samples^108^ (Table S2) were consolidated across samples into single chromosome GenomicsDB workspaces via GATK GenomicsDBImport^136^. Each workspace was joint-genotyped with GATK GenotypeGVCFs^137^, and the resulting chromosome population VCFs were combined using the GatherVcfs function in the Picard toolkit^135^. The joint-genotyped population VCF was processed and filtered with bcftools version 1.13^134^ following the guidelines published by the Haploteam for population joint-genotyping *S. cerevisiae*^138^. The published protocol is based on the GATK best practices for identifying germline short variants^139^. The only parameters that were modified were a) added a minimum quality threshold of 30 b) minimum genotype quality was increased from 20 to 30, c) the initial genotype missingness threshold was lowered from 99% to 80%, d) excess heterozygosity was set to 0.95. The final joint call set contained 2,950 samples that passed filters.

### Determination of Cell Ploidy

The DNA content of the sequenced strains (a measure of ploidy level) was determined by staining cells with SYTOX Green Nucleic Acid Stain (Thermo Fisher Scientific). The preparation of cells for DNA content analysis was performed as follows. For each sample, 5 × 10L cells were collected, washed with ddH₂O, resuspended in 400 µL ddH₂O, and sonicated. Then, 950 µL of 100% ethanol was added to the sample, and the mixture was stored at −20°C overnight to fix the cells. Cells were collected by centrifugation, resuspended in 800 µL of 50 mM sodium citrate containing 250 µg/mL ribonuclease A (Thermo Fisher Scientific), and incubated at 50°C overnight. Next, 50 µL of 20 mg/mL proteinase K (Invitrogen) was added to each sample, followed by incubation at 50°C for 2 hours. Cells were collected by centrifugation, resuspended in 50 mM sodium citrate containing 2 µM SYTOX Green Nucleic Acid Stain (Thermo Fisher Scientific), and incubated at room temperature for 2 hours, protected from light. Cells were briefly sonicated to reduce clumping and analyzed by flow cytometry. SYTOX Green intensity was measured using a 488 nm laser and a 536/40 (FITC) band-pass filter on a Sysmex CyFlow. The fluorescent signal of previously established haploid (CSH377), diploid (CSH556), and tetraploid (CSH369) strains were used to generate a calibration curve. Genome-wide ploidy and chromosomal aneuploidy profiles were further validated computationally with AStra^140^ using the sorted BAM files (Figure S10, Table S10).

### SNP-based Phylogenomic Inference

The population genotyped VCF was further filtered with bcftools to only retain SNP variant sites, for a SNP gap of 10 bases, and to only retain sites with a minimum genotyping rate of at least 99%. The resulting VCF contained 1,877,031 genome-wide SNP sites and 2,950 samples. The VCF was converted to FASTA format using the script vcf2phylip.py^141^, with heterozygous genotypes randomly resolved. Maximum-likelihood-like tree inference was performed with VeryFastTree 4.0^142,143^ under the GTR nucleotide model. Node supports were estimated with 1000 single-likelihood-branch test bootstrap replicates (Figure S1). The resulting phylogeny was visualized in iTOL version 7.5^144^ and rooted with the CHN-IX/TW1 lineage as an outgroup. Branches were colored according to broad evolutionary groups. Populations were assigned through identification of monophyletic lineages, retaining populations previously identified in the literature when possible. In addition, a subsampled tree restricted to 209 strains isolated from baking environments (Table S10) was extracted from the full phylogeny to serve a backbone for visualizing ploidy, aneuploidy, and heterozygosity of baking-associated strains. Similarly, a subtree was pruned for 111 isolates for which maltose metabolism growth assays were performed (Table S11) and for 43 isolates for which sporulation efficiency and spore viability were measured (Table S12).

### Population Structure Analyses

The VCF containing 2,950 samples employed in phylogenetic reconstruction (Table S2) was further filtered for examination of *S. cerevisiae* population structure with both model-free and model-based methods. PLINK version 1.9^145,146^ was used to remove sites with any missing data, remove samples with more than 5% missing genotypes, select only biallelic SNPs, and remove variants with a minor allele frequency lower than 5%. Variants were thinned to retain 1 SNP every 1000 base pairs to ensure independence. SNP thinning by windows was done instead of traditional pruning for linkage disequilibrium because *S. cerevisiae* can reproduce asexually and by selfing, so filtering for linkage disequilibrium with near-clones present in the dataset would remove a majority of the variants. After filtering, 2,853 samples and 5,274 sites remained and were converted to PLINK binary BED format. Principal components analysis was conducted in PLINK 1.9^145,146^ with default parameters and 20 principal components were retained. The R statistical computing environment version 4.5.0^147^ was used to produce a scree plot to assess the percentage of variance explained by each principal component (Figure S4) and to generate 2D scatter plots for every combination of principal components to visualize population structure (Figure S5). Plots were made with the R package ggplot2^148^. FastMixture version 1.3.0^149^ was run on the same PLINK bed file for K values 2-40 with the *–cv=10* option enabled to perform 10-fold cross-validation. Cross-validation errors for each K value were plotted with the R package ggplot2^148^ (Figure S2). The base R stats package^147,150^ and the segmented package^151^ were used to fit a piecewise linear regression model to the cross-validation errors for each value of K, and breakpoint analysis was used to determine the optimal K (number of hypothetical ancestral populations). The first point at which the slope between two K values was no longer significantly different from zero was selected as the best K value (K=6) (Figure S2, Figure 2B). Q-matrices of inferred ancestry proportion were plotted as stacked bar plots for all values of K (Figure S3).

### Population Summary Statistics

Genome-wide nucleotide diversity (pi), pairwise nucleotide difference (*D_xy_*) and pairwise *F_ST_* were calculated for broad evolutionary groups and finer-scale populations corresponding to the global phylogeny. Calculations for pi and *D_xy_* (Table S4-5) were done with Pixy version 1.25^152,153^ and pairwise *F_ST_* (Table S3) was calculated with Piawka version 0.8.11^154^. The summary statistics were calculated from an all-site genotype VCF with minimal filtering (GATK Best Practices site filtration, no INDELs, minimum depth 10, and maximum 20% missingness) to prevent biasing calculations through the exclusion of invariant sites. The proportion of heterozygosity was estimated for each newly sequenced sample using an in-house script (Github/Zenodo Link) (Table S10). A heatmap for population pairwise *F_ST_* was produced with the R package ggalign^155^.

### Pangenome Analysis

Pangenome gene content and copy number were examined through a “map-to-pan” approach. Short reads for the 40 newly sequenced isolates, along with the 1,333 publicly available isolates previously mapped to the *S. cerevisiae* reference genome, were aligned to a recently published *S. cerevisiae* gene-based pangenome reference from Leogler et al.^111^ The pangenome gene-based reference contained 8,541 contigs, each representing a pangenome gene sequence. Short reads were aligned to the reference FASTA file using BWA-mem2^156^ with the options *-t 20 -U 0 -L 0,0 -O 4,4*. Reads were sorted with Samtools version 1.13^134^, read groups were assigned with GATK AddOrReplaceReadGroups^136^, and then alignments were indexed with Samtools. Normalized pangenome gene copy number was calculated as per Loegler et al.^111^ by adapting their published scrips. The parameters *-x 0.15 -t 0.3* were used for read depth normalization and gene presence detection thresholds respectively. Methods and parameters for read alignment, read processing, and downstream gene copy number calling strictly adhered to those employed by Loegler et al. to ensure compatible integration with previously published pangenome gene copy number profiles for an additional 1,610 isolates^111^. The pangenome gene copy number profiles for each sample were combined to form a large pangenome matrix. A binary gene presence/absence matrix was created from the normalized copy number matrix in R version 4.5.0^147^, setting any value under 0.3 to absent (0), and any value over 0.3 to present (1). Any samples with more than 80% absent genes were removed from the dataset, as those likely had low sequencing depth/coverage or low mapping quality. Any genes that were not present in a single sample were removed. The final datasets consisted of two matrices, one of copy number and one of presence/absence, for 2,913 strains and 8,535 genes (Data S1, Data S2).

The pangenome PAV dataset was used to categorize genes as core (present in all samples), softcore (present in at least 95% of samples), distributed (present in less than 95% of samples, but more than 1 sample), and private (only present in a single sample) (Table S7, Figures S6-S7). A heatmap of presence/absence variation (PAV) was generated for the distributed genes across samples from the six broad evolutionary groups using the R package ggalign^155^. The full pangenome gene presence/absence profiles for 2,913 strains were converted to FASTA format by collapsing the matrix columns into a single sequence per strain and used as input to IQ-TREE version 2.3^157^ to infer a pangenome gene PAV phylogeny. Phylogenetic inference was performed under the binary GTR2 model with ultrafast bootstrap approximation^158^ and the options *-safe -czb -B 1000 -nm 5000*. The resulting maximum-likelihood phylogeny was visualized and annotated in iTOL version 7.5^144^ (Figure S8). Genome size (as a function of number of pangenome genes) was calculated for each sample by counting the genes present in the pangenome PAV matrix (Table S8). Genome size was plotted by broad evolutionary groups and populations determined through phylogenetic and population structure analyses using the ggplot2^148^ R package. A Kruskal-Wallis test and post-hoc pairwise Dunn Test was conducted in R version 4.5.0^147,150^ to test for significant differences in the number of pangenome genes among populations (Table S9).

### Maltose Growth Assay

Strains (Table S11) were inoculated in 96-well microtiter plates in 200 µL of SC media (per liter 1.7 g yeast nitrogen base without amino acid and without ammonium sulfate, 5 g ammonium sulfate, 2 g amino acid mix, 2% dextrose) and incubated at 30°C with shaking overnight. The overnight cultures were back diluted to an OD600 of 0.1-0.2 in 200 µL of SC+maltose media (per liter 1.7 g yeast nitrogen base without amino acid and without ammonium sulfate, 5 g ammonium sulfate, 2 g amino acid mix, 2% maltose). Growth was measured using an Epoch2 Microplate Reader (Biotek Instruments) with OD600 measured every 15 minutes with continuous orbital shaking at 30°C over 48 hours. The OD600 readings were extracted from the Epoch2 Microplate Reader files and the time was converted to hours. The OD600 data was modeled with the Growthcurver R package^159^ and the values for the initial population size (N_0_), the maximum population size (K), the intrinsic growth rate of the population (r), and the populations doubling time (t_gen) were calculated. Per-strain outliers were removed from growth rate data by calculating z-scores for all replicates and removing those with z-scores below -2 and above 2. A sub-tree was pruned from the global SNP-based phylogeny for the 111 strains included in the maltose growth assay and maltose growth was plotted along the phylogeny using the R packages ggplot2^148^, ggtree^160^, and ggtreeExtra^161^. A list of maltose-associated genes was obtained from the Saccharomyces Genome Database^131^ and combined with novel genes detected in the pangenome reference that were annotated with likely maltose-associated functions^111^. Copy number for the maltose-associated genes was extracted from the pangenome gene copy number dataset for each of the strains phenotyped for maltose growth (Data S3). A copy number heatmap for maltose-associated pangenome genes was aligned to the maltose strain subtree and the maltose growth rate plot using ggtreeExtra^161^. Presence/absence profiles for the maltose-associated genes were used to count the total number detected in each strain. Linear regression and phylogenetically independent spearman rank correlation analyses were performed in R version 4.5.0^147^ with the stats^150^ and MultiTraits^162^ R packages to test the correlation between maltose growth rate and quantity of maltose associated genes, as well as between maltose growth rate and copy number of each maltose-associated gene (Figure S11).

### Sporulation and Spore Viability

*S. cerevisiae* strains (Table S12) were first streaked out onto YPD plates and grown for ∼48Lh at 30 °C. A single colony was cultured overnight in 5 mL of liquid YPD (1% yeast extract, 2% peptone, 2% dextrose) at 30 °C with three biological replicates per strain. The following morning, each culture was transferred to a 15 mL conical tube and centrifuged at 2000 rpm for 2Lmin. Cell pellets were resuspended in 450 μL ddH2O, and 150 μL of each mixture was spread onto solid SPM plates (1% potassium acetate, 0.1% yeast extract, 2% agar, 0.05% dextrose) and left to sporulate at 25°C for 7 days. After this 7-day incubation, a quarter of the lawn of each plate was harvested using a bent pipette tip and suspended in 1LmL ddH2O. Cells were counted on a hemocytometer, and percent sporulated cells calculated with 2 technical replicates per biological replicate (Table S12). To calculate spore viability, strains were sporulated as above. Sporulated cells were resuspended in a mixture of 25 μL ddH2O and 2LμL of 5 mg/mL 20LT Zymolyase (the equivalent of 1 U). This suspension was incubated in a 37L°C water bath for 10 min, then 500 μL of ddH2O was added to halt the Zymolyase reaction. Tetrads were dissected on YPD plates using a Singer SporePlay+ microscope (Singer Instruments). Spore viability was calculated as the number of spores that successfully returned to a vegetative state divided by the total number of spores dissected. 4 tetrads (16 total possible colonies) were analyzed per strain (Table S10). A sub-tree was pruned from the global SNP-based phylogeny for the 43 strains included in the sporulation efficiency and spore viability assays and served as the y-axis to plot sporulation results (Figure S12).

### Computation, Statistical Analysis, and Visualization

Bioinformatic pipelines and comparative genomic analyses were executed on the North Carolina State University Bioinformatics Resource Center high performance computing cluster. Statistical analyses and visualizations were executed in R version 4.5.0 unless otherwise stated.

### Data availability

Genome sequencing data generated from this study are available at NCBI SRA PRJNA1446738. Strains isolated by the Heil lab (Table S1) are available upon request.

## ACKNOWLEDGEMENTS

We thank Caiti Lahue, Jessica McNeill, and Nico Sevier for assistance in data collection. We appreciate comments on this manuscript from members of the Heil Lab past and present. We acknowledge members of the Dunn Lab, Aminah Al-Attas Bradford, Hanna Berman, Bradley Alff, and Sam Niewierowski, for fruitful discussions and feedback. We are grateful to all the researchers from the Global Sourdough Project (Elizabeth Landis, Angela Oliverio, Erin McKenney, Lauren Nichols, Nicole Kfoury, Megan Biango-Daniels, Leonora Shell, Anne Madden, Lori Shapiro, Shravya Sakunala, Kinsey Drake, Albert Robbat, Matthew Booker, and Noah Fierer) and to the bakers who donated their sourdough starters. Some strains used in phenotypic assays are courtesy of Joseph Schacherer. This work was funded by NIH R35 GM142849 to C.S.H.

## Notes

### Competing Interest Statement

The authors have declared no competing interest.

