## Supplemental Figures for "Insights from sourdough redefine the domestication landscape of baker’s yeast"

Margot Ruffieux[1], Nathan Brandt[1], Alexxis Guttierez[1,2], Benjamin E. Wolfe[3], Rob R. Dunn[4], Caiti Smukowski Heil[1*]

Affiliations:

1 Department of Biological Sciences, North Carolina State University, Raleigh, NC
2 Current address: Saint Louis University School of Medicine, St. Louis, MO
3 Department of Biology, Tufts University, Medford, MA
4 Department of Applied Ecology, North Carolina State University, Raleigh, NC

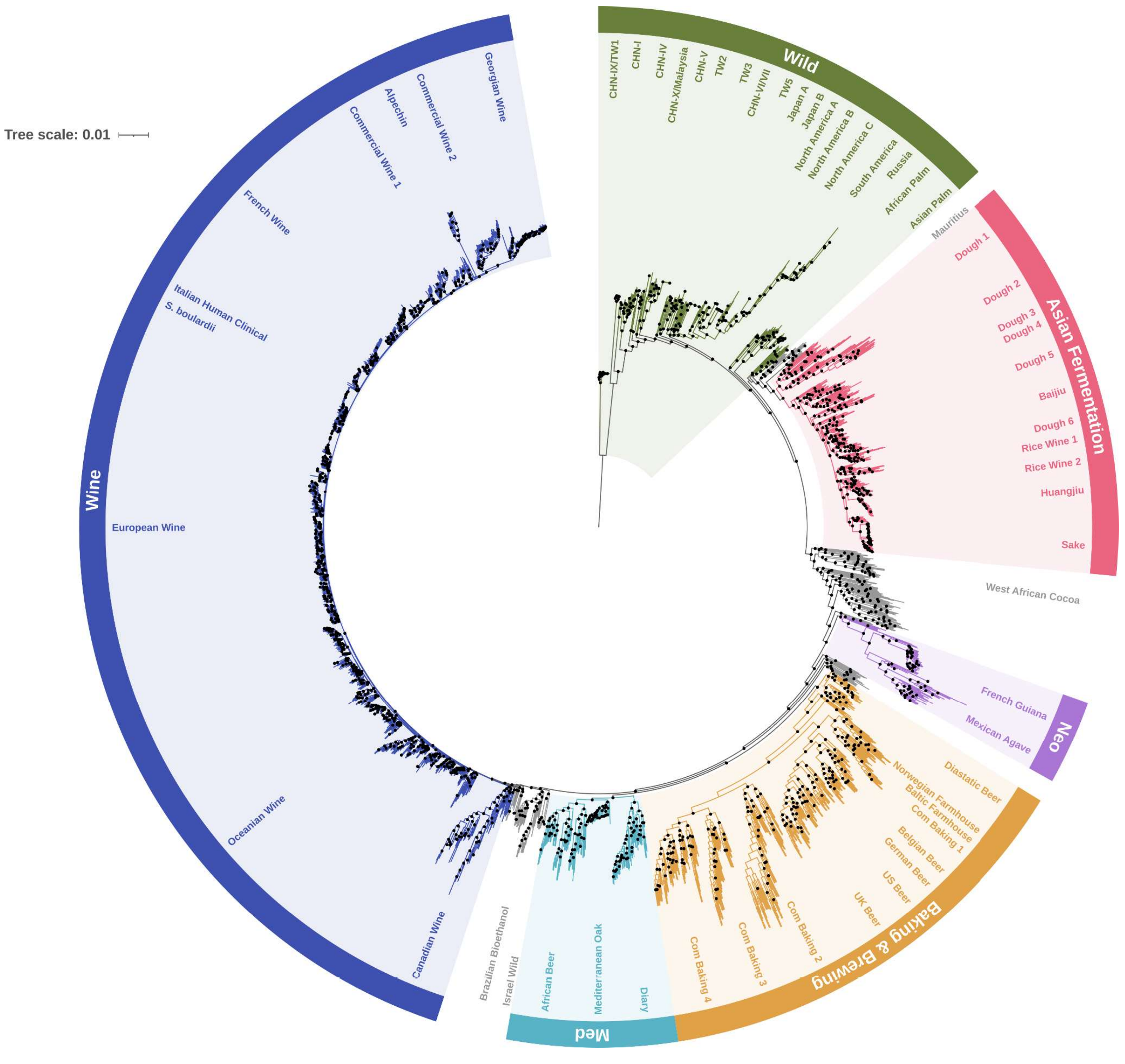


Figure S1. SNP-based maximum-likelihood-like phylogeny with black circles indicating nodes with bootstrap support values 90% and above.

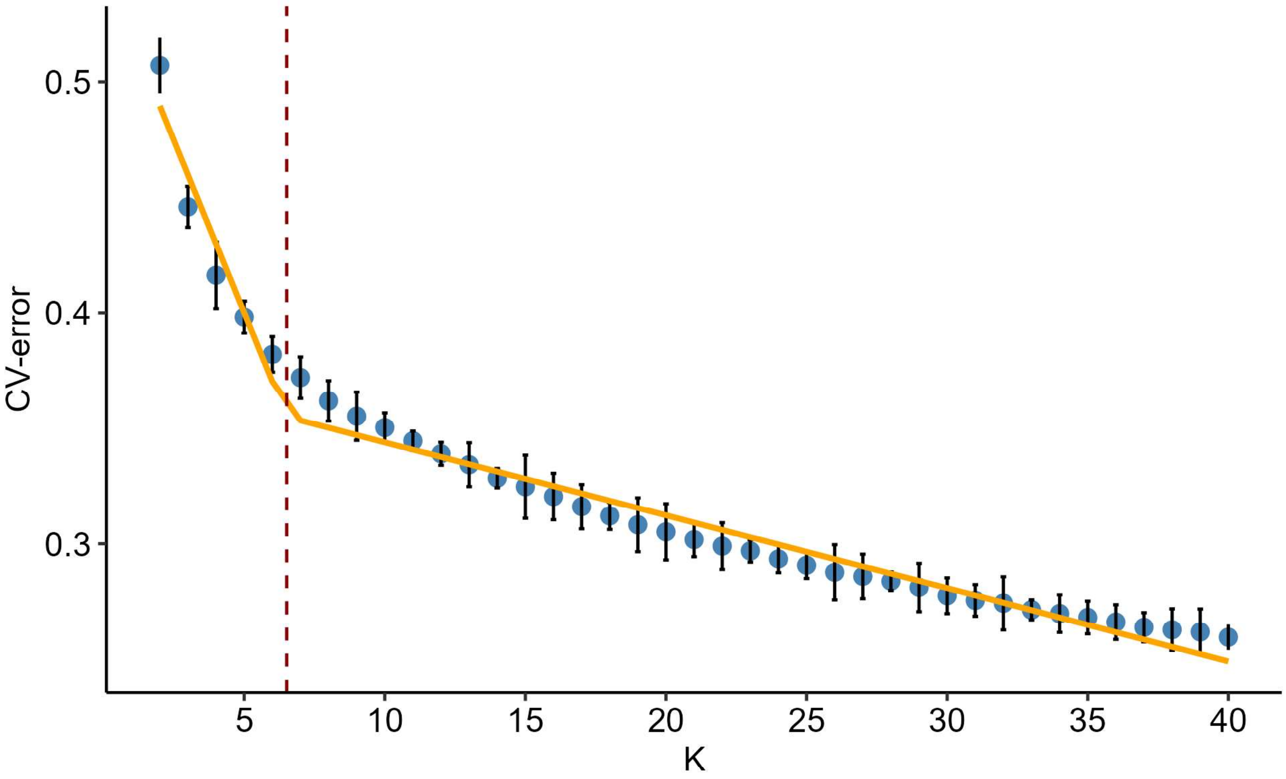


Figure S2. FastMixture ten-fold cross-validation errors and breakpoint regression analysis for determining optimal value of K.

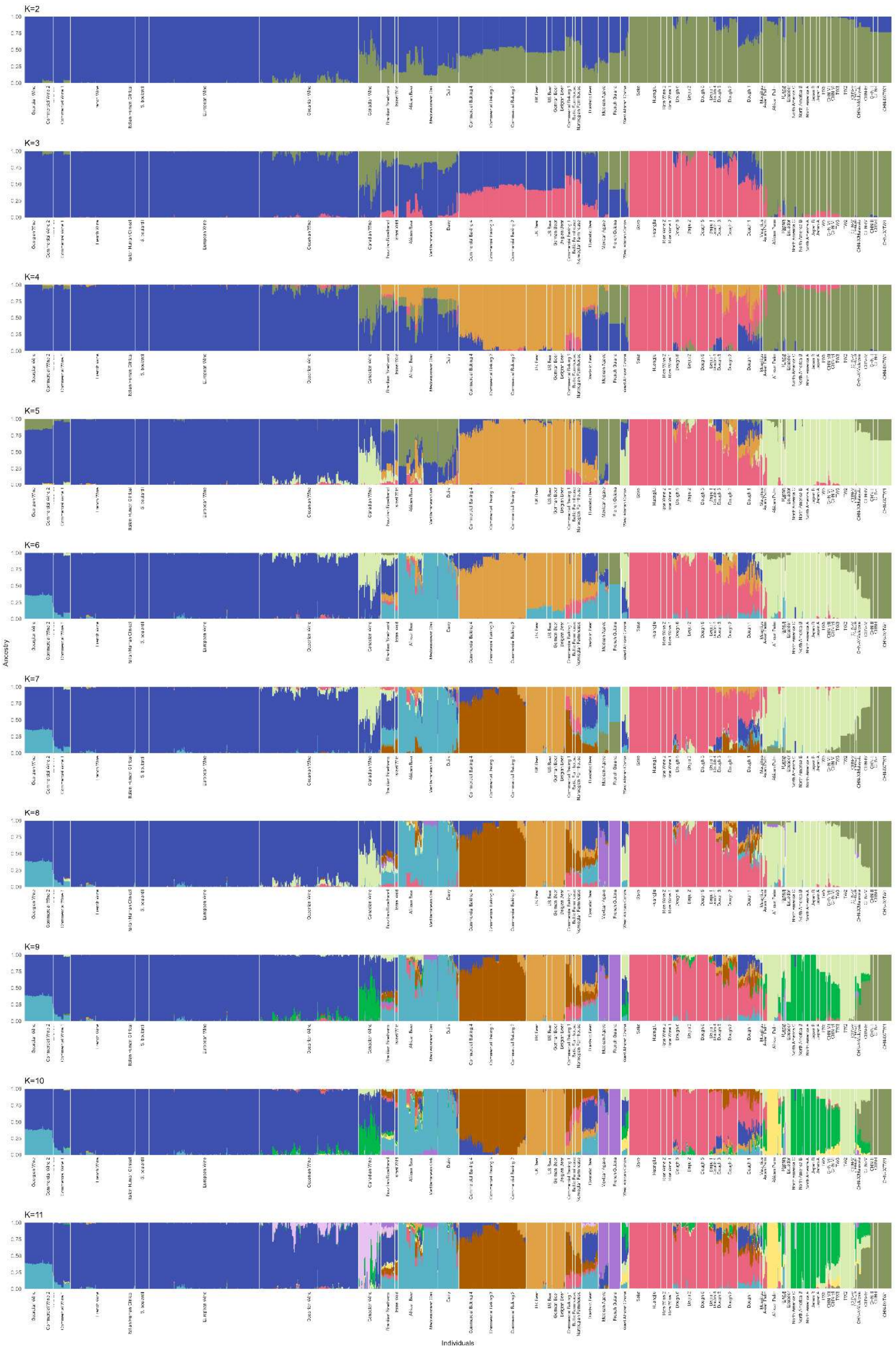

K=2
K=3
K=4
K=5
K=6
K=7
K=8
K=9
K=10
K=11
Ancestry
Individuals

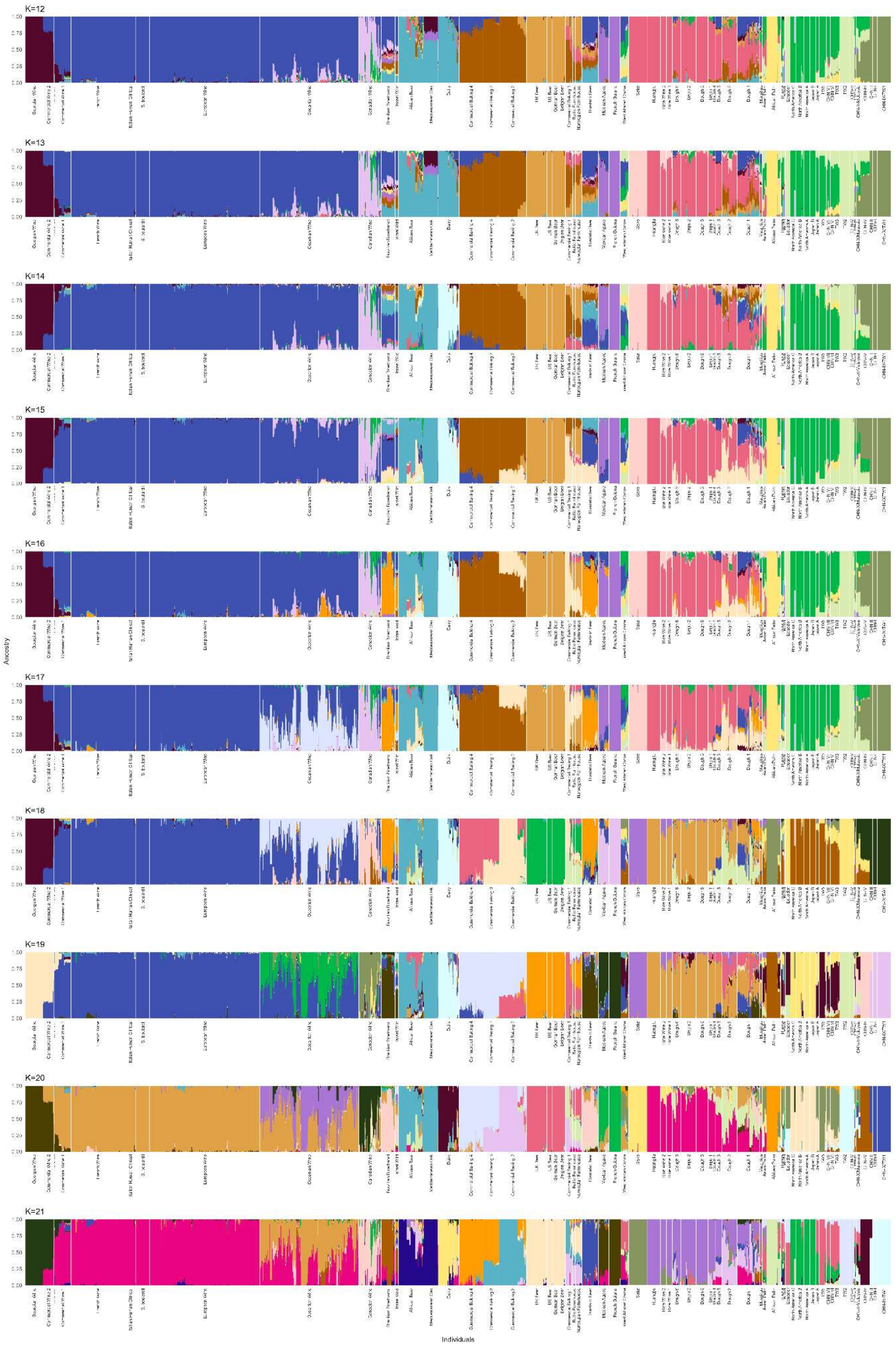
K=12
K=13
K=14
K=15
K=16
K=17
K=18
K=19
K=20
K=21
Ancestry
Individuals

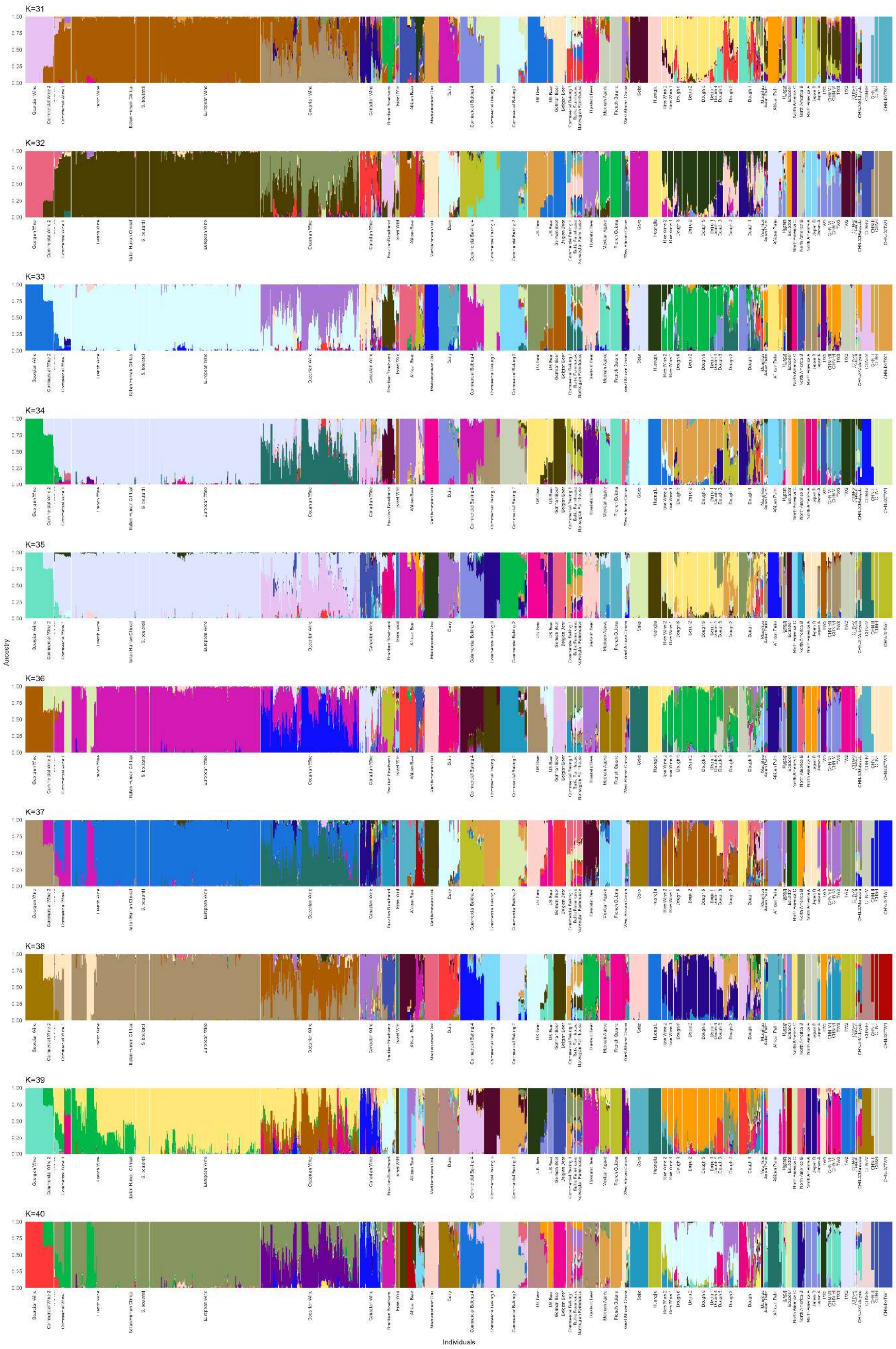
K=31
K=32
K=33
K=34
K=35
K=36
K=37
K=38
K=39
K=40
Ancestry
Individuals

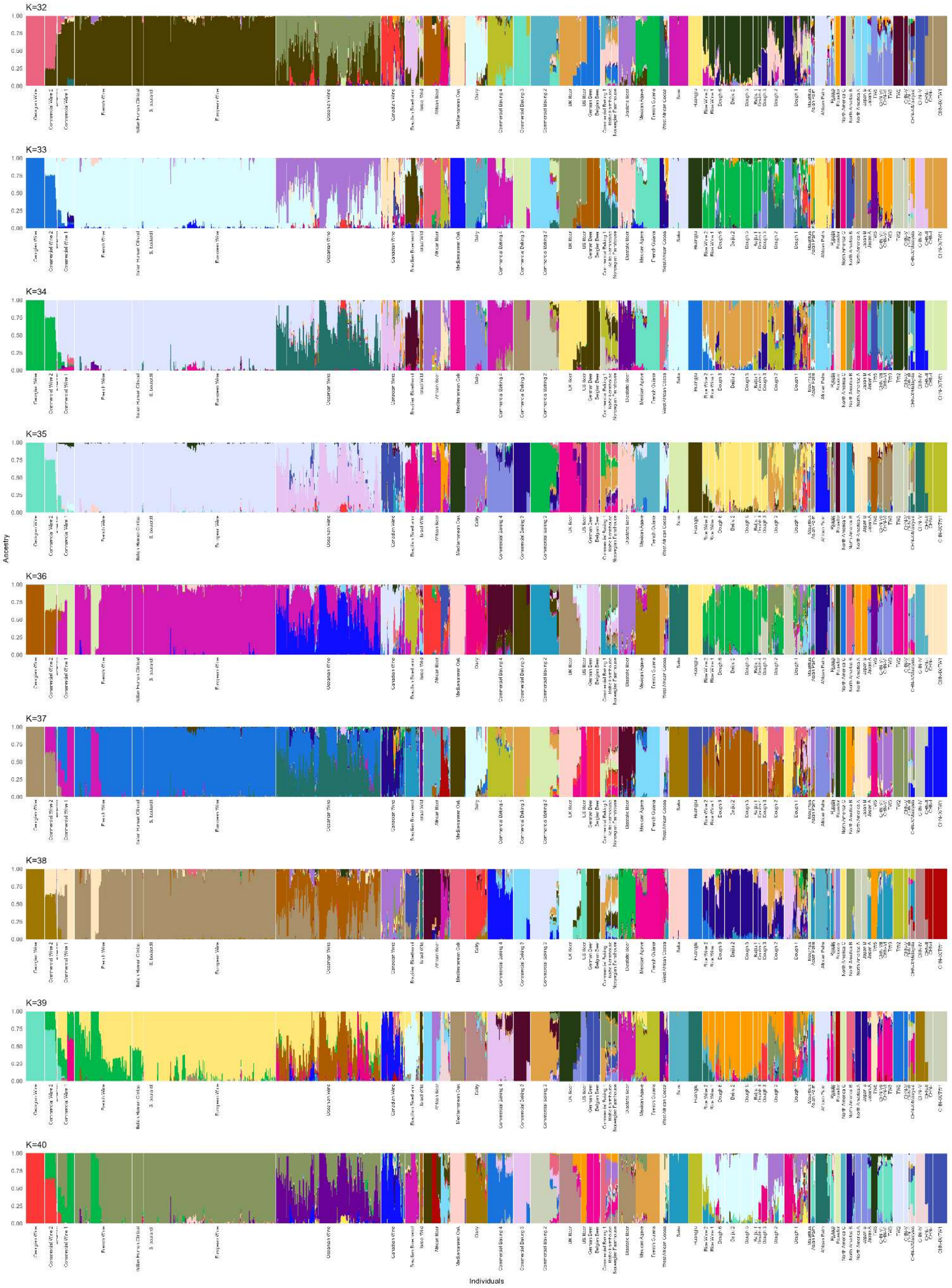


Figure S3. Stacked bar plots of estimated ancestry proportion from FastMixture analysis for K values 2 through 40.

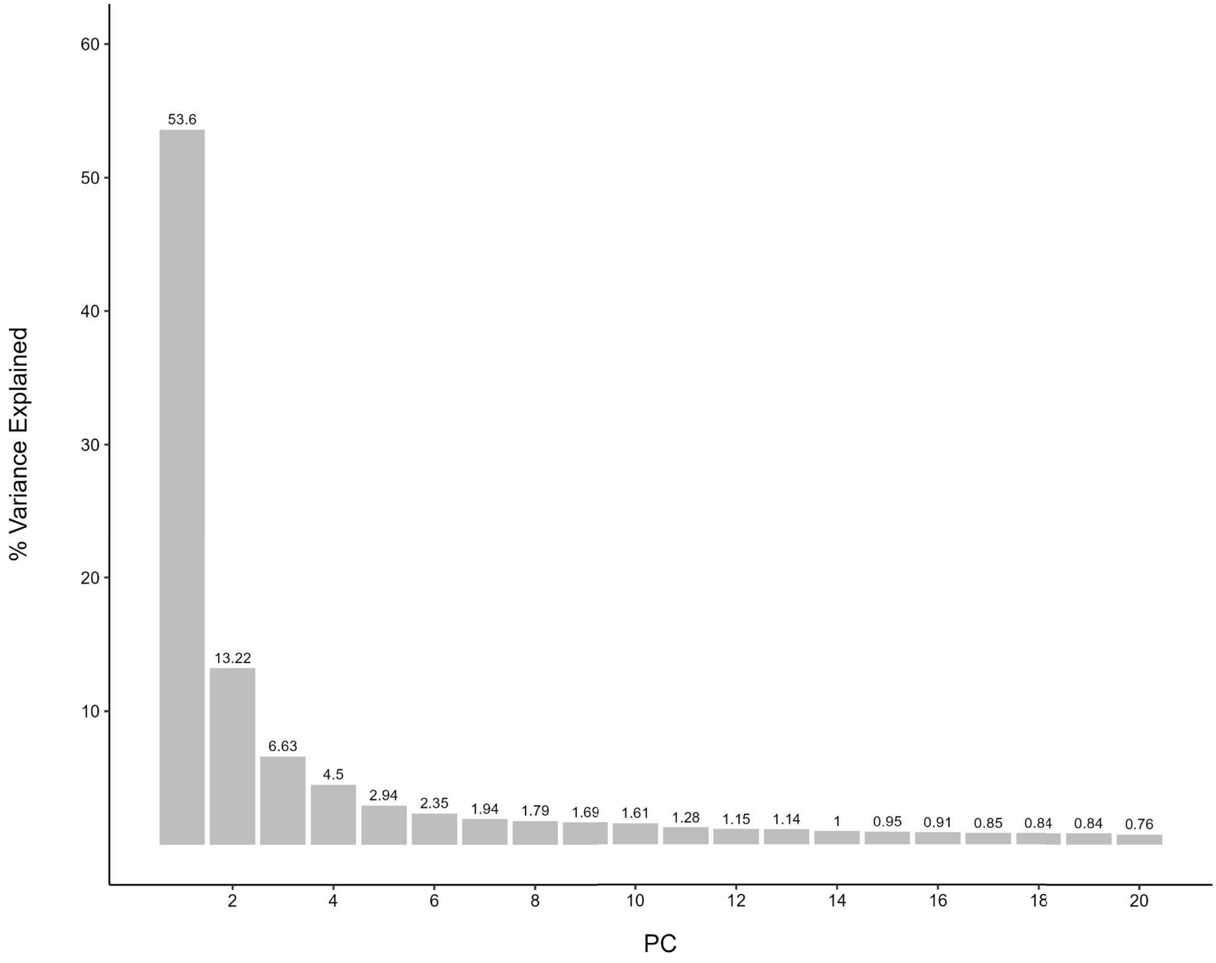


Figure S4. Scree plot depicting percentage of variance explained by the first 20 principal components obtained through Principal Component Analysis.

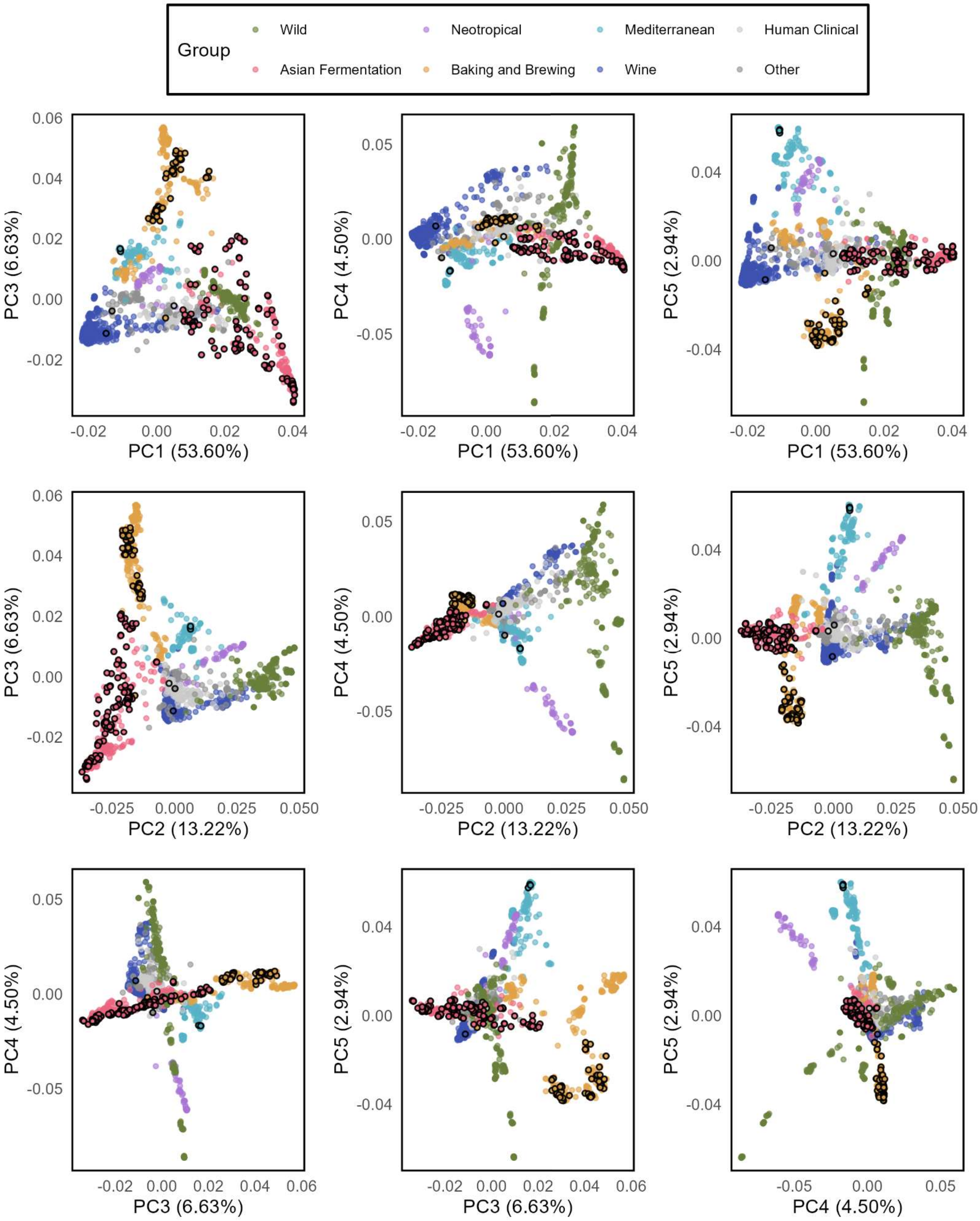


Figure S5. Scatter plots of each combination of Principal Components for PC1 through PC6.

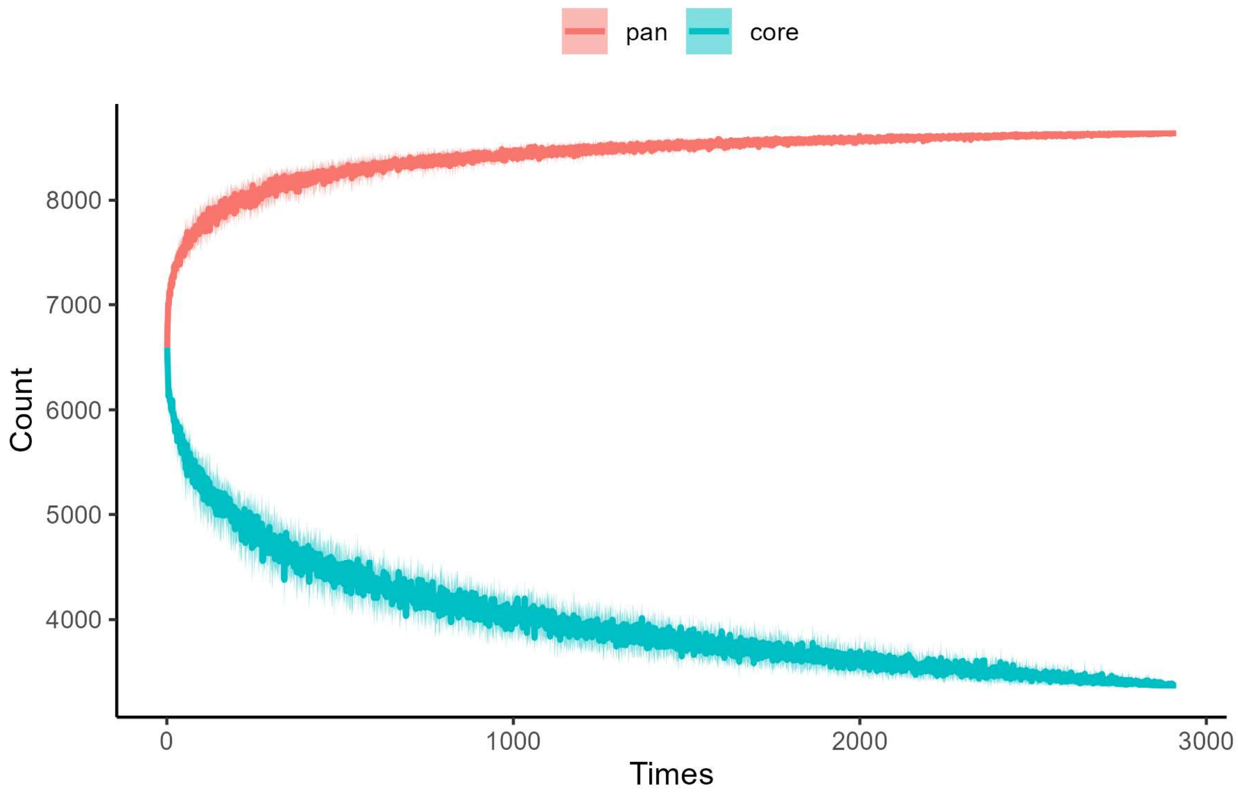


Figure S6. Gene accumulation curve simulated for core and pangenome genes.

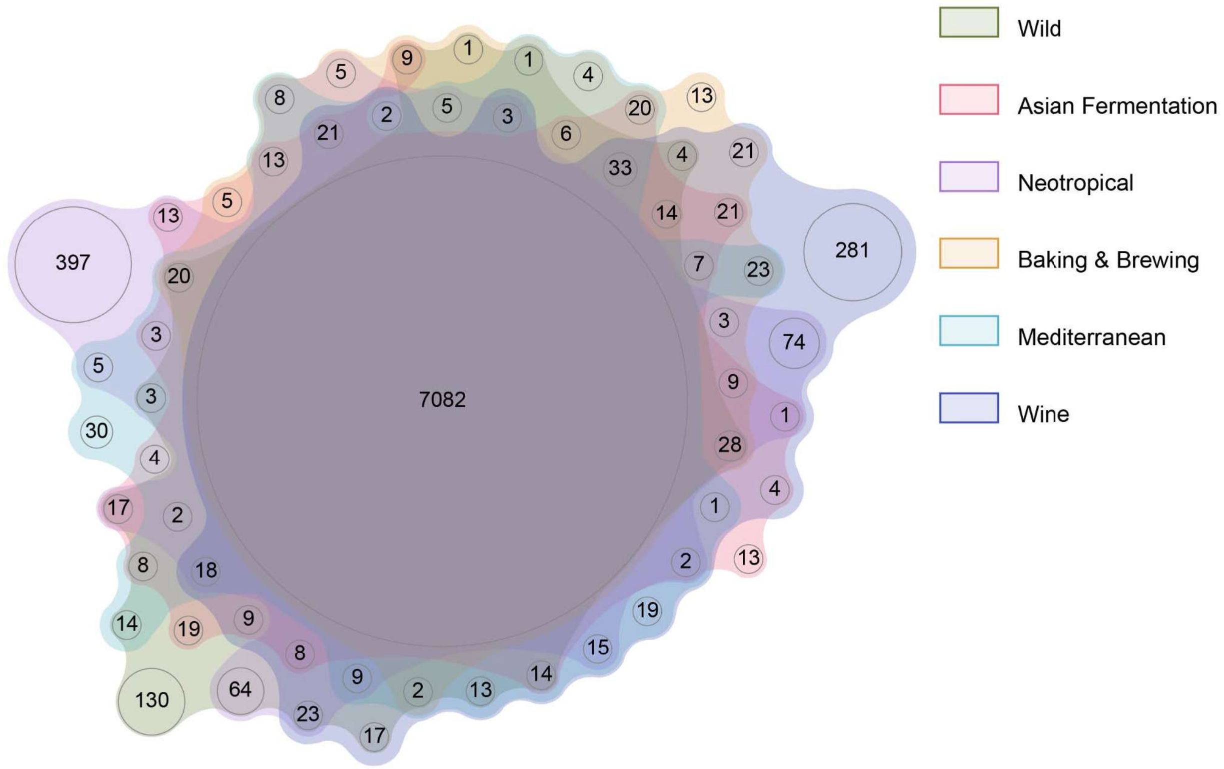


Figure S7. Venn diagram of six-way intersection for pangenome gene PAVs of six major evolutionary groups.

A

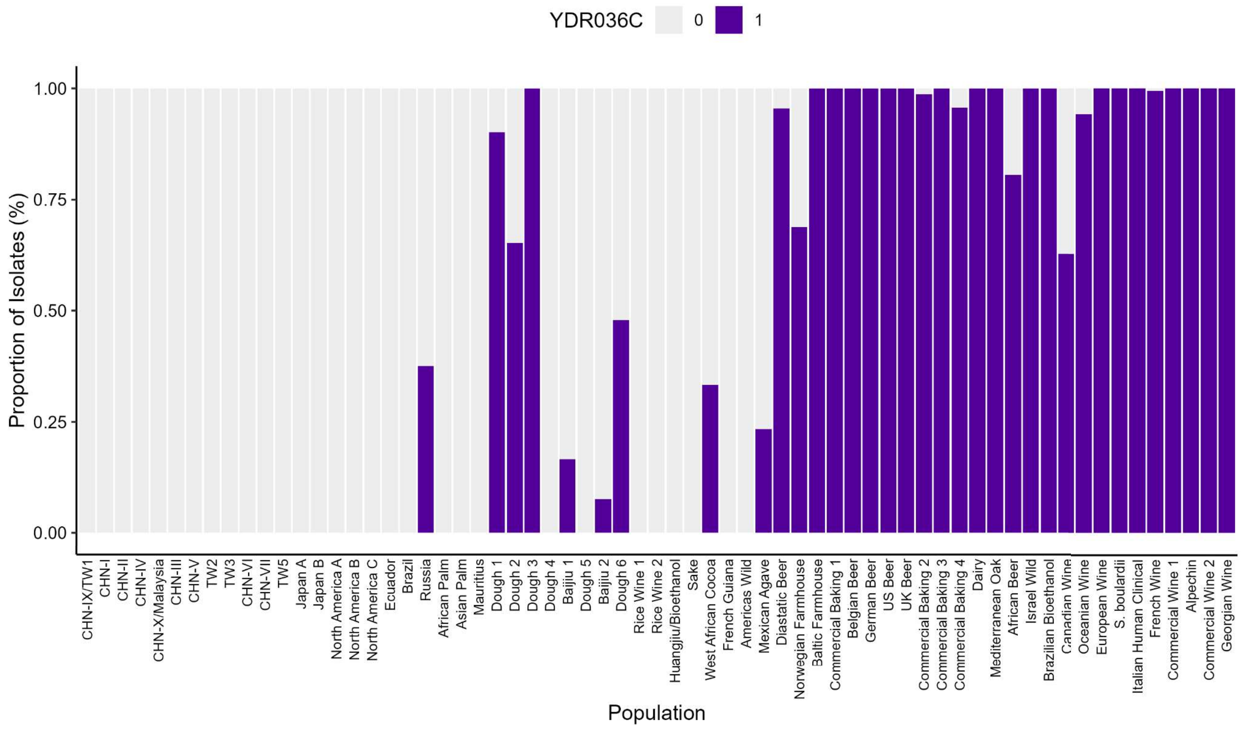
YDR036C
0
1
Proportion of Isolates (%)
1.00
0.75
0.50
0.25
0.00
Population
CHN-IX/TW1
CHN-I
CHN-II
CHN-IV
CHN-X/Malaysia
CHN-III
CHN-V
TW2
TW3
CHN-VI
CHN-VII
TW5
Japan A
Japan B
North America A
North America B
North America C
Ecuador
Brazil
Russia
African Palm
Asian Palm
Mauritius
Dough 1
Dough 2
Dough 3
Dough 4
Baijiu 1
Dough 5
Baijiu 2
Dough 6
Rice Wine 1
Rice Wine 2
Huangjiu/Bioethanol
Sake
West African Cocoa
French Guiana
Americas Wild
Mexican Agave
Diastatic Beer
Norwegian Farmhouse
Baltic Farmhouse
Commercial Baking 1
Belgian Beer
German Beer
US Beer
UK Beer
Commercial Baking 2
Commercial Baking 3
Commercial Baking 4
Dairy
Mediterranean Oak
African Beer
Israel Wild
Brazilian Bioethanol
Canadian Wine
Oceanian Wine
European Wine
S. boulardii
Italian Human Clinical
French Wine
Commercial Wine 1
Alpechin
Commercial Wine 2
Georgian Wine

B

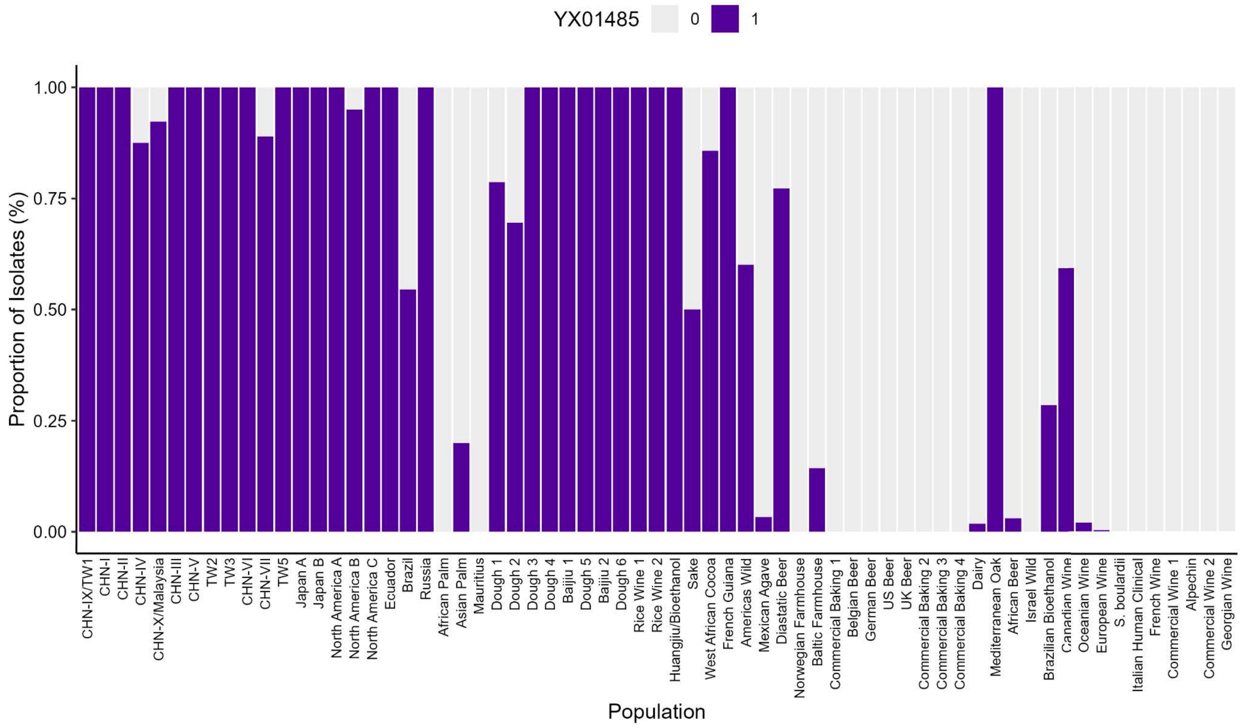
YX01485
0
1
Proportion of Isolates (%)
1.00
0.75
0.50
0.25
0.00
Population
CHN-IX/TW1
CHN-I
CHN-II
CHN-IV
CHN-X/Malaysia
CHN-III
CHN-V
TW2
TW3
CHN-VI
CHN-VII
TW5
Japan A
Japan B
North America A
North America B
North America C
Ecuador
Brazil
Russia
African Palm
Asian Palm
Mauritius
Dough 1
Dough 2
Dough 3
Dough 4
Baijiu 1
Dough 5
Baijiu 2
Dough 6
Rice Wine 1
Rice Wine 2
Huangjiu/Bioethanol
Sake
West African Cocoa
French Guiana
Americas Wild
Mexican Agave
Diastatic Beer
Norwegian Farmhouse
Baltic Farmhouse
Commercial Baking 1
Belgian Beer
German Beer
US Beer
UK Beer
Commercial Baking 2
Commercial Baking 3
Commercial Baking 4
Dairy
Mediterranean Oak
African Beer
Israel Wild
Brazilian Bioethanol
Canadian Wine
Oceanian Wine
European Wine
S. boulardii
Italian Human Clinical
French Wine
Commercial Wine 1
Alpechin
Commercial Wine 2
Georgian Wine

C

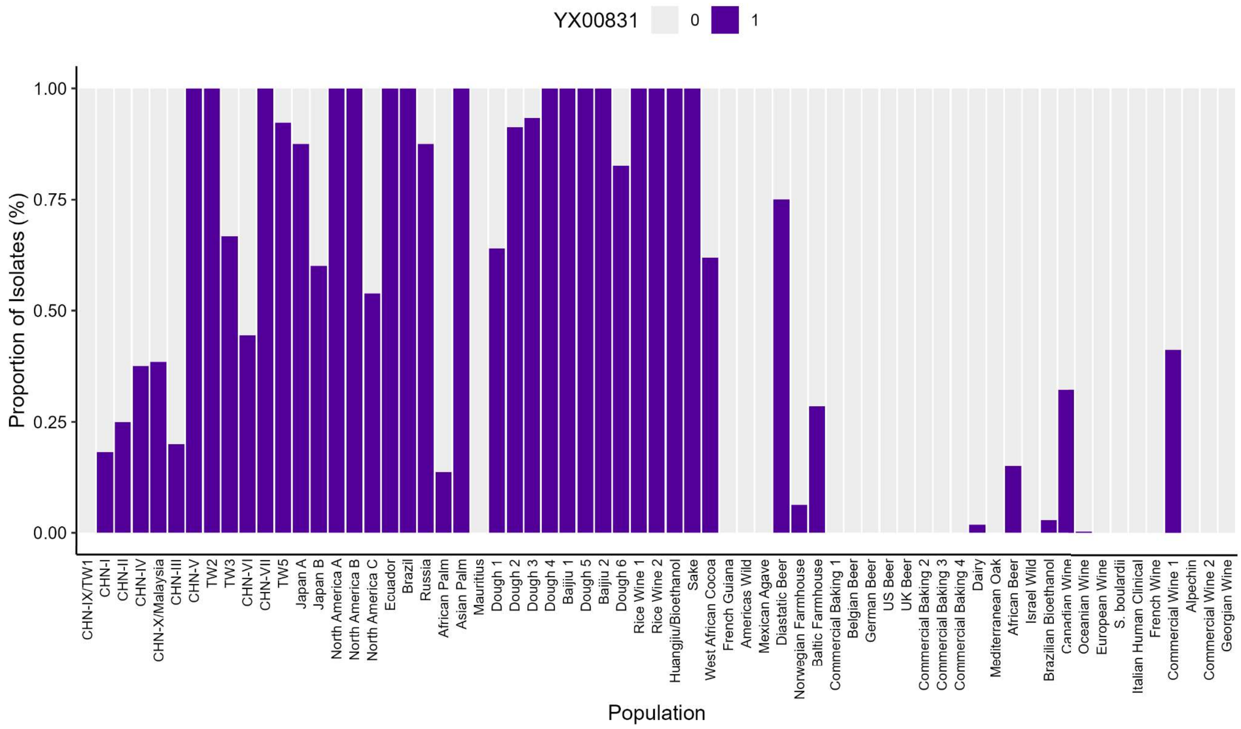
YX00831
0
1
Proportion of Isolates (%)
1.00
0.75
0.50
0.25
0.00
Population
CHN-IX/TW1
CHN-I
CHN-II
CHN-IV
CHN-X/Malaysia
CHN-III
CHN-V
TW2
TW3
CHN-VI
CHN-VII
TW5
Japan A
Japan B
North America A
North America B
North America C
Ecuador
Brazil
Russia
African Palm
Asian Palm
Mauritius
Dough 1
Dough 2
Dough 3
Dough 4
Baijiu 1
Dough 5
Baijiu 2
Dough 6
Rice Wine 1
Rice Wine 2
Huangjiu/Bioethanol
Sake
West African Cocoa
French Guiana
Americas Wild
Mexican Agave
Diastatic Beer
Norwegian Farmhouse
Baltic Farmhouse
Commercial Baking 1
Belgian Beer
German Beer
US Beer
UK Beer
Commercial Baking 2
Commercial Baking 3
Commercial Baking 4
Dairy
Mediterranean Oak
African Beer
Israel Wild
Brazilian Bioethanol
Canadian Wine
Oceanian Wine
European Wine
S. boulardii
Italian Human Clinical
French Wine
Commercial Wine 1
Alpechin
Commercial Wine 2
Georgian Wine

D

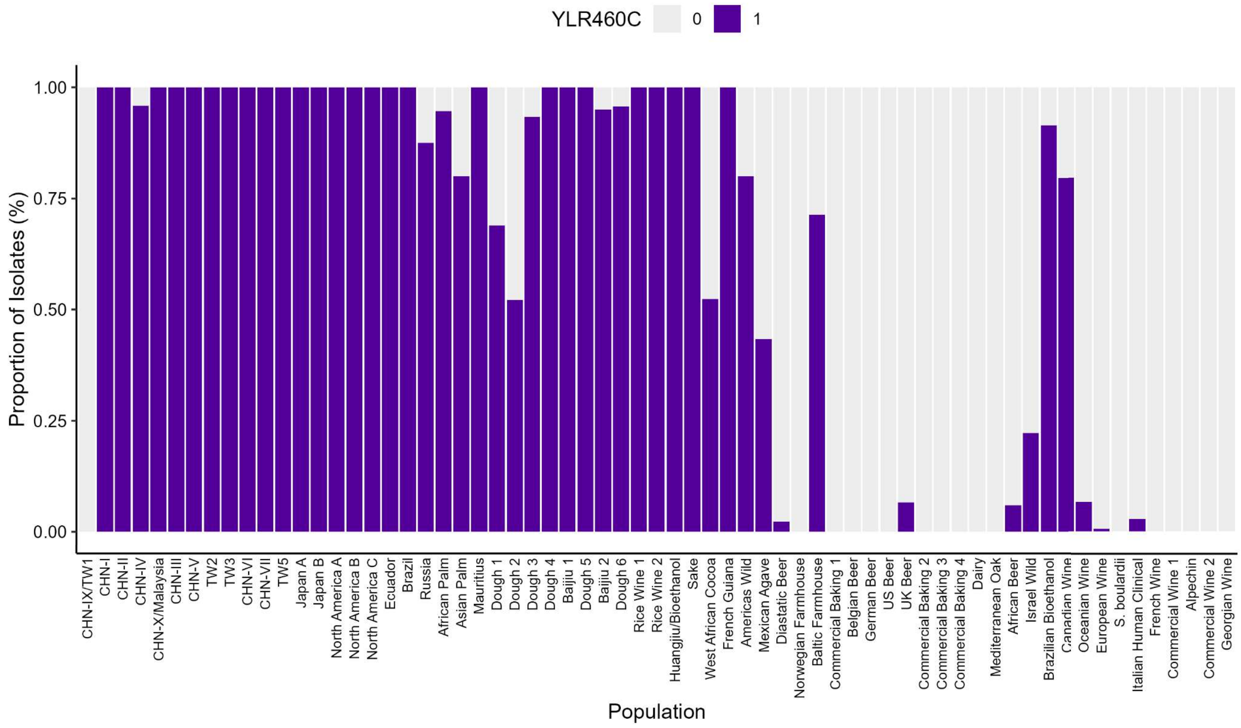
YLR460C
0
1
Proportion of Isolates (%)
1.00
0.75
0.50
0.25
0.00
Population
CHN-IX/TW1
CHN-I
CHN-II
CHN-IV
CHN-X/Malaysia
CHN-III
CHN-V
TW2
TW3
CHN-VI
CHN-VII
TW5
Japan A
Japan B
North America A
North America B
North America C
Ecuador
Brazil
Russia
African Palm
Asian Palm
Mauritius
Dough 1
Dough 2
Dough 3
Dough 4
Baijiu 1
Dough 5
Baijiu 2
Dough 6
Rice Wine 1
Rice Wine 2
Huangjiu/Bioethanol
Sake
West African Cocoa
French Guiana
Americas Wild
Mexican Agave
Diastatic Beer
Norwegian Farmhouse
Baltic Farmhouse
Commercial Baking 1
Belgian Beer
German Beer
US Beer
UK Beer
Commercial Baking 2
Commercial Baking 3
Commercial Baking 4
Dairy
Mediterranean Oak
African Beer
Israel Wild
Brazilian Bioethanol
Canadian Wine
Oceanian Wine
European Wine
S. boulardii
Italian Human Clinical
French Wine
Commercial Wine 1
Alpechin
Commercial Wine 2
Georgian Wine

E

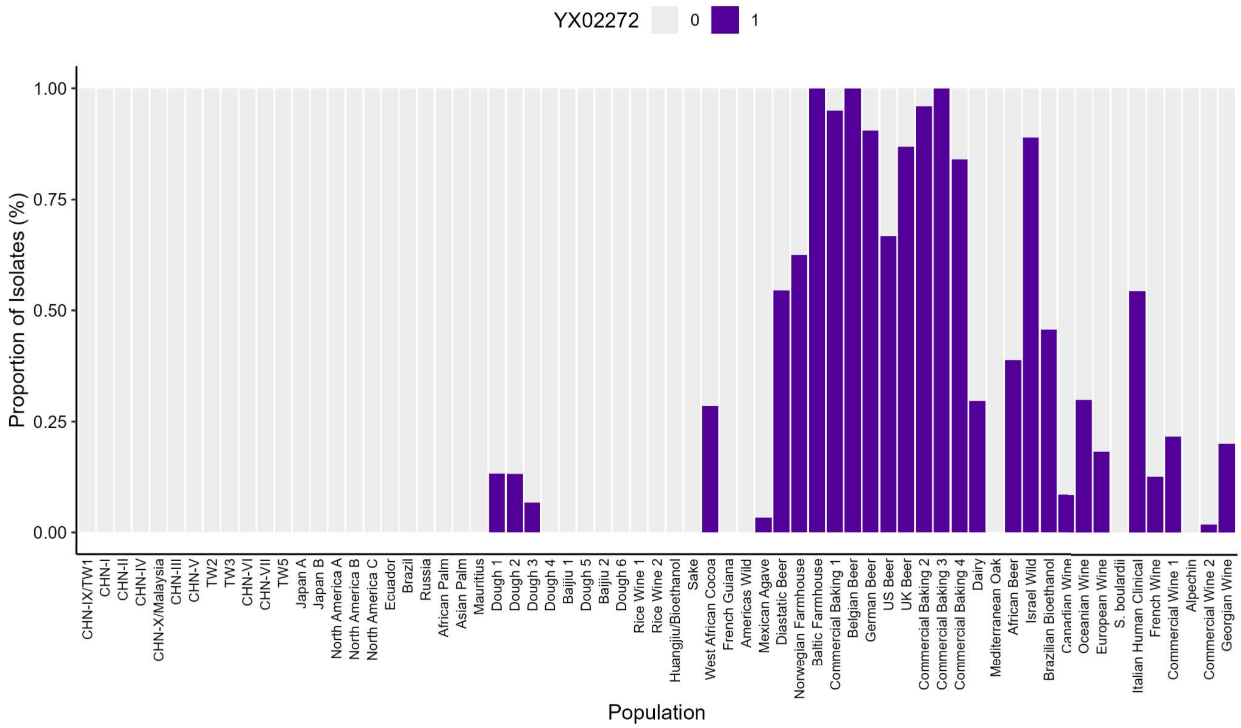


Figure S8. Frequencies of selected representative pangenome genes in *S. cerevisiae* populations. A) YDR036 gene separates west Eurasian from east Eurasian domestication trajectories. B-D) YX01845, YX00831, and YLR460C genes differentiate east Eurasian from west Eurasian domestication trajectories. E) YX02272 is predominately found in beer and commercial baking populations.

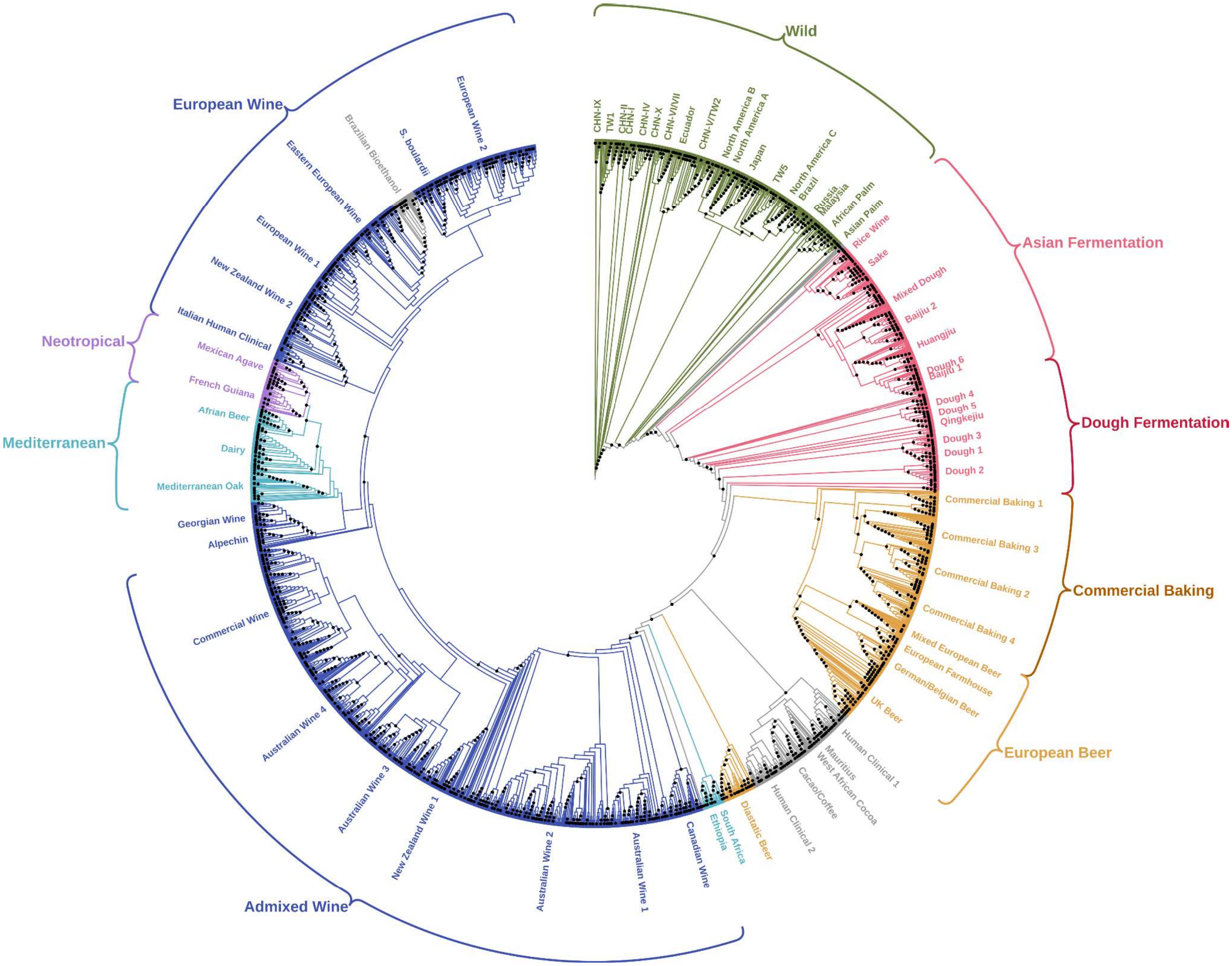


Figure S9. Pangenome binary gene presence/absence phylogeny with nodes supported by bootstrap values at or above 90% annotated by black circles.

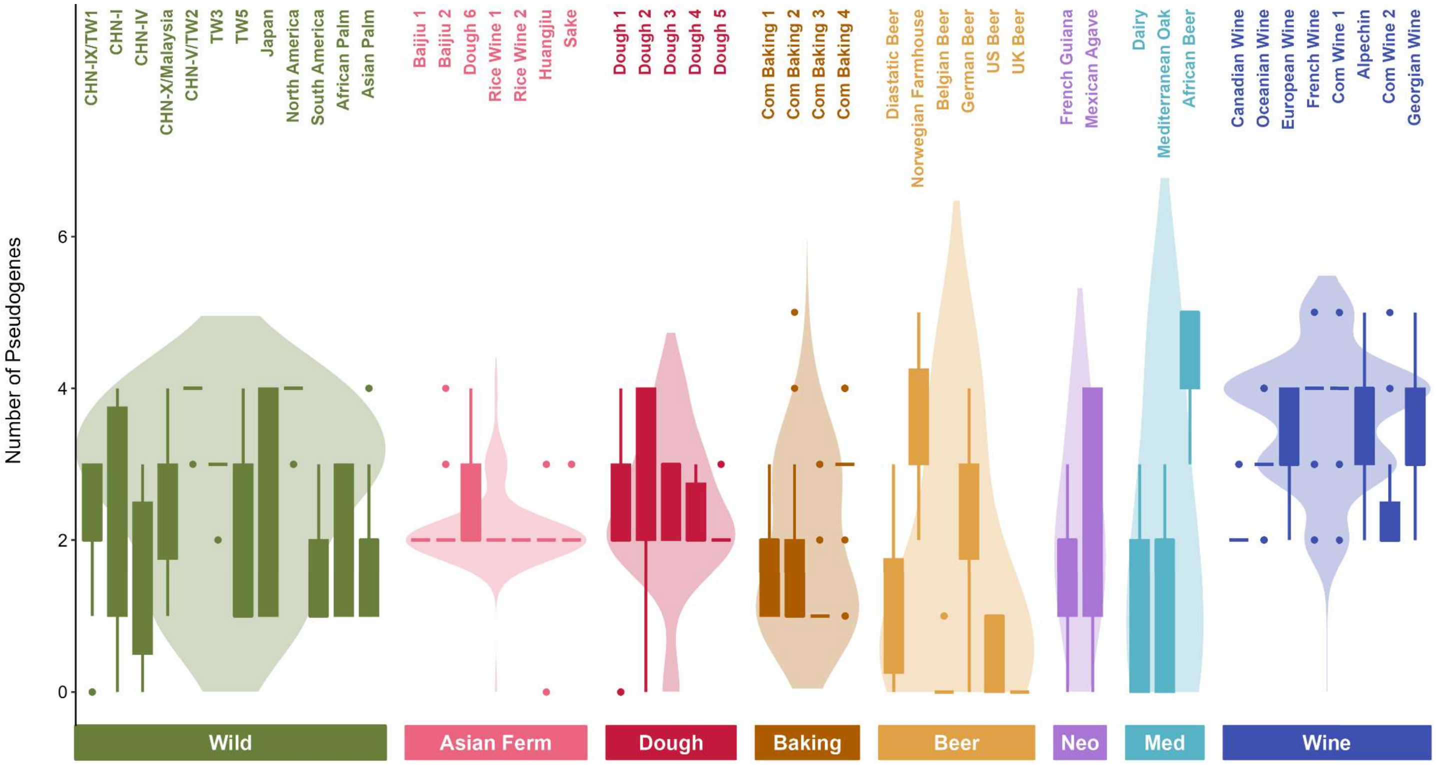


Figure S10. Number of pseudogenes per strain by population and broad evolutionary group.

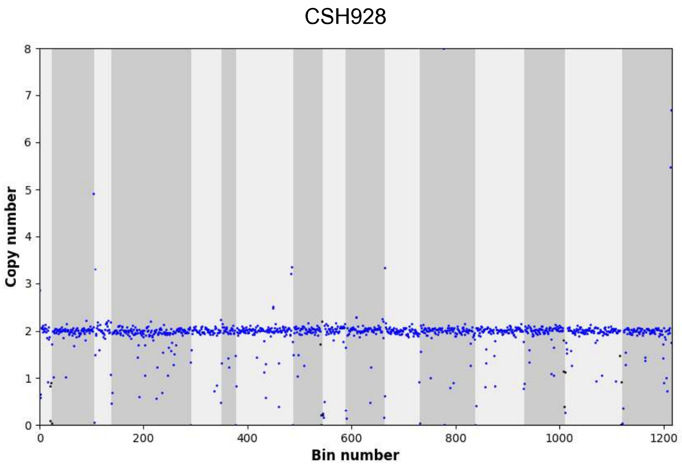
CSH928
Copy number
Bin number

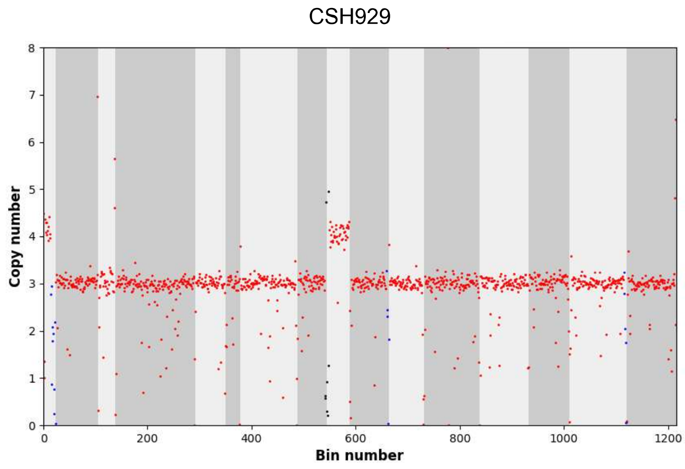
CSH929
Copy number
Bin number

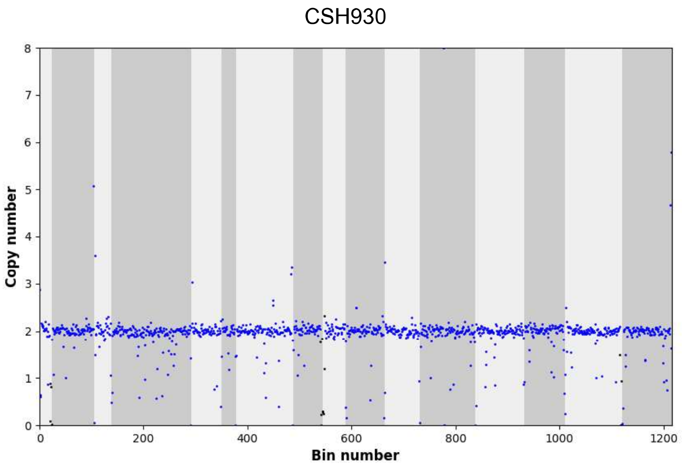
CSH930
Copy number
Bin number

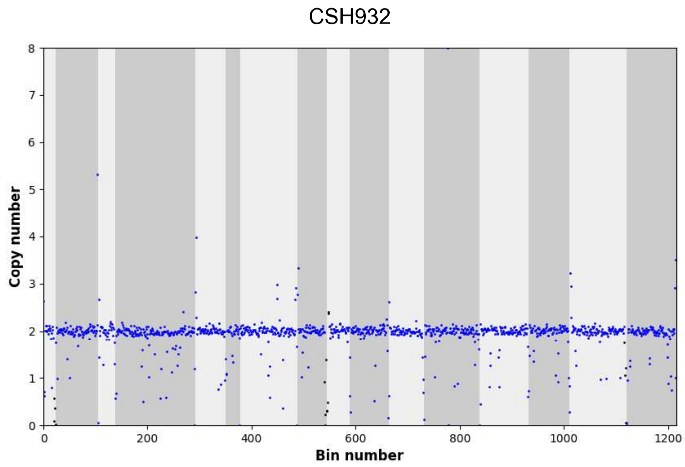
CSH932
Copy number
Bin number

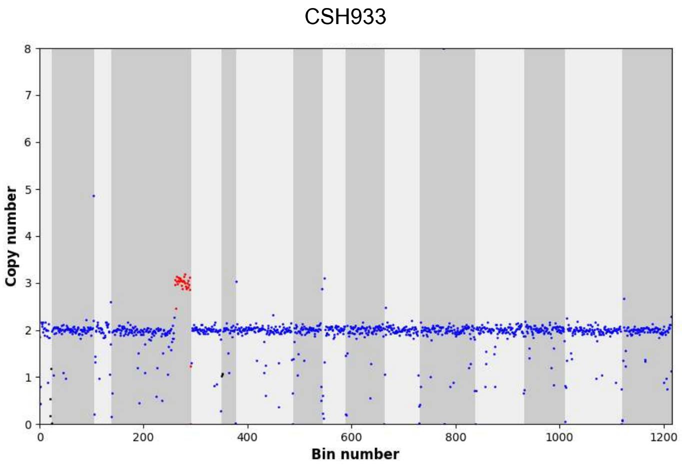
CSH933
Copy number
Bin number

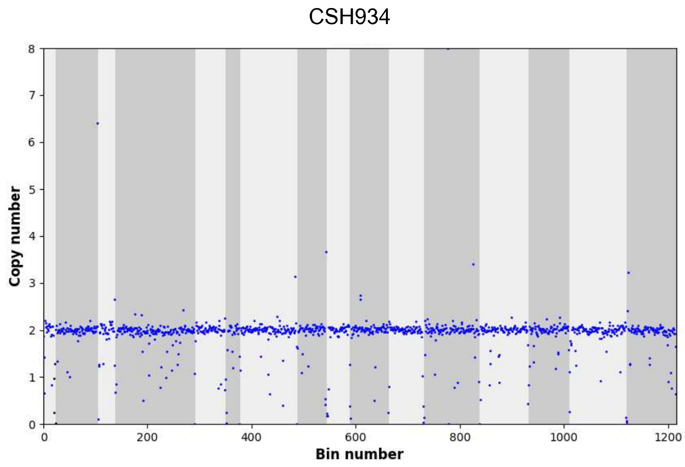
CSH934
Copy number
Bin number

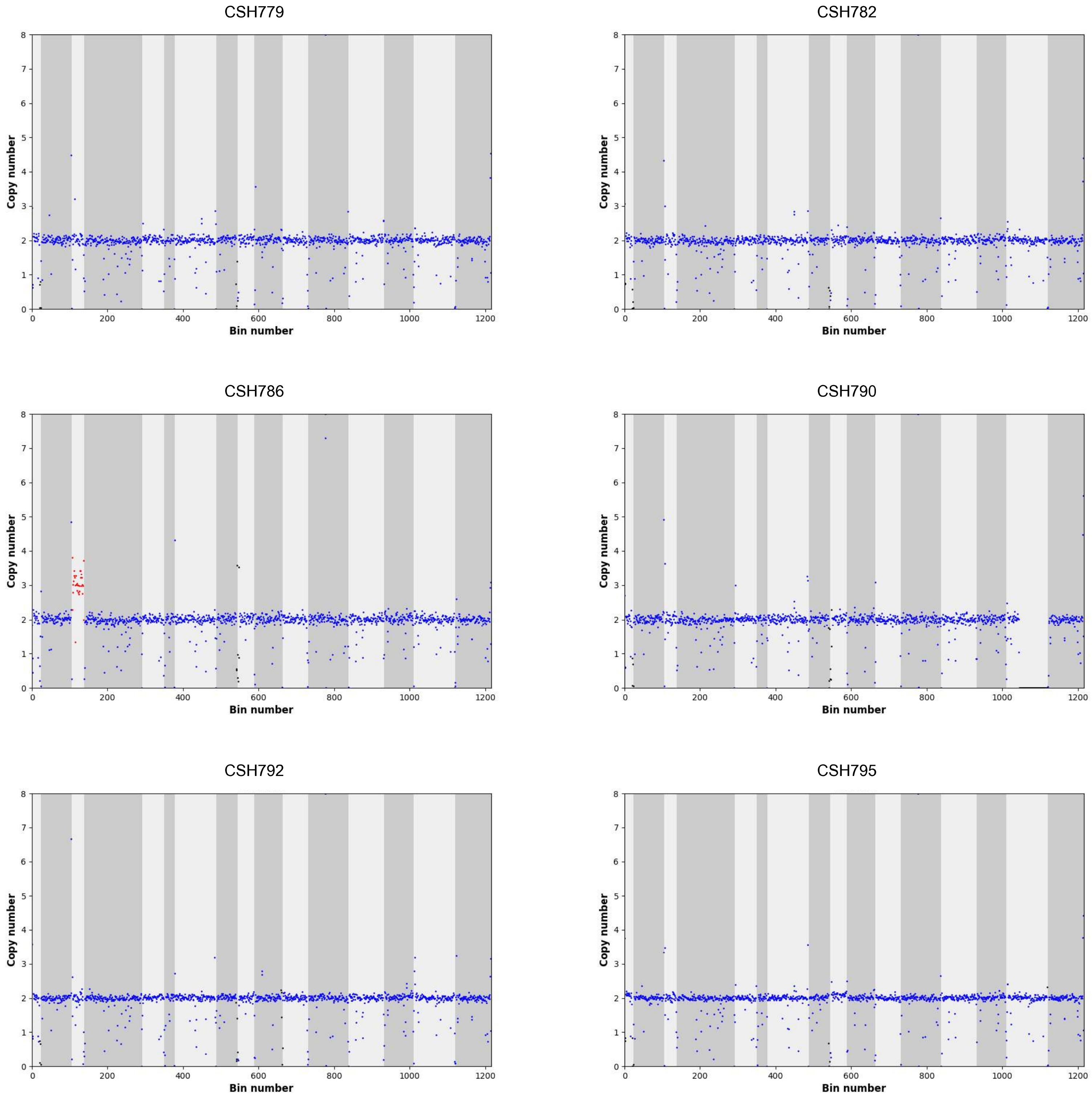
CSH779
CSH782
CSH786
CSH790
CSH792
CSH795
Copy number
Bin number
0
1
2
3
4
5
6
7
8
200
400
600
800
1000
1200

CSH808

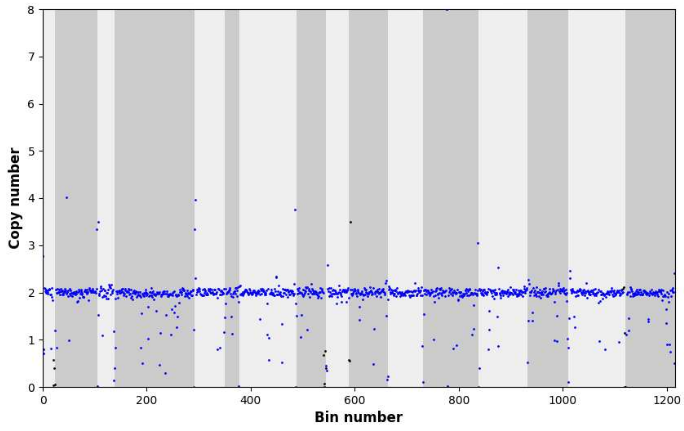


CSH811

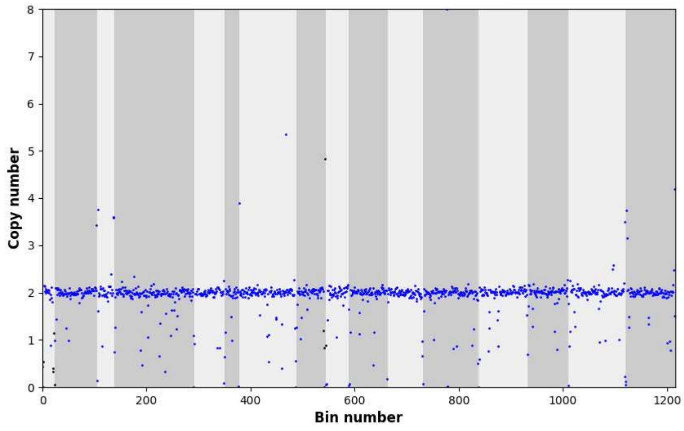


CSH816

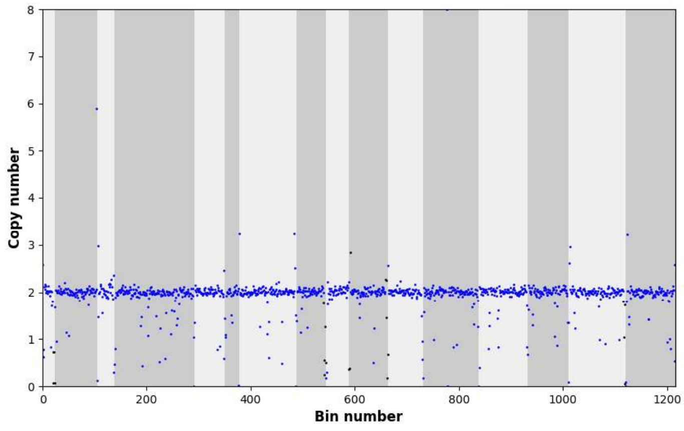


CSH819

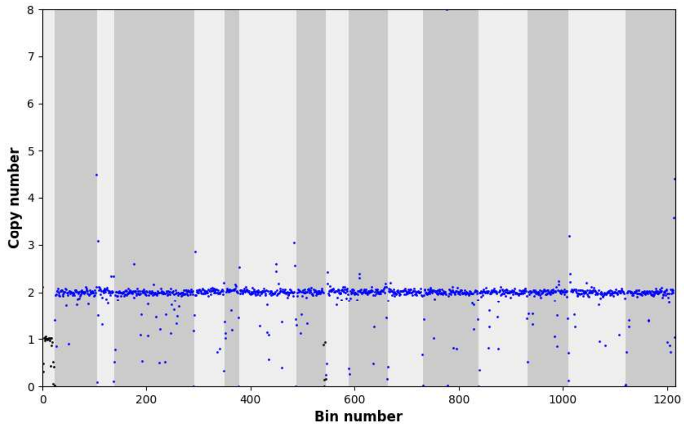


CSH823

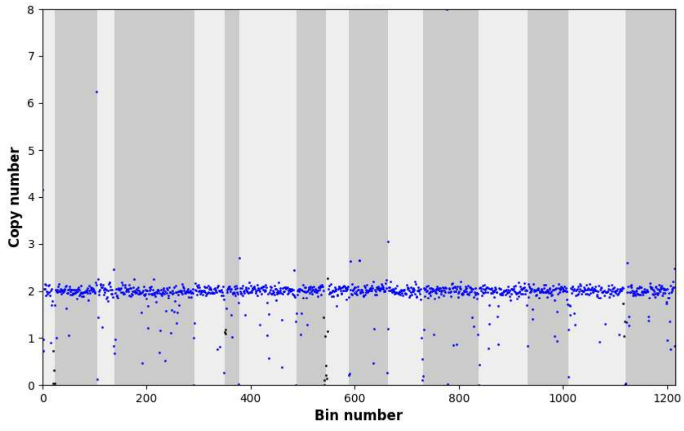


CSH824

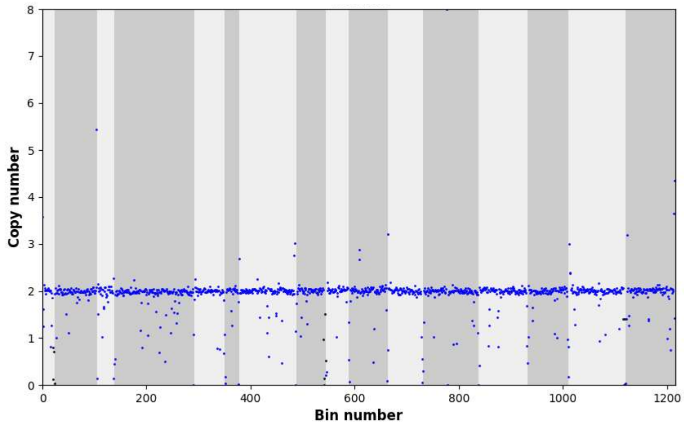

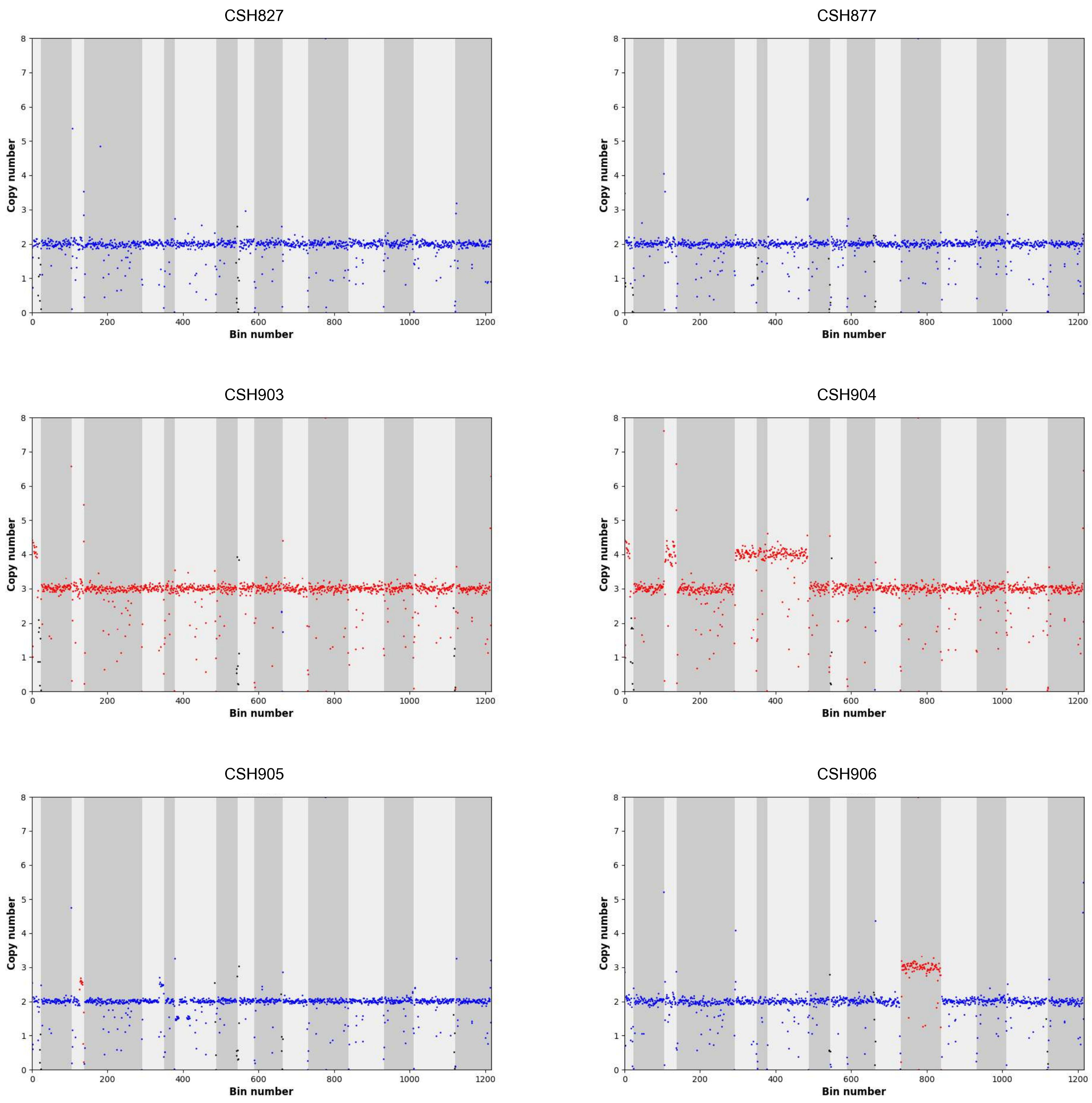
CSH827
CSH877
CSH903
CSH904
CSH905
CSH906
Copy number
Bin number
0 1 2 3 4 5 6 7 8
0 200 400 600 800 1000 1200

CSH907

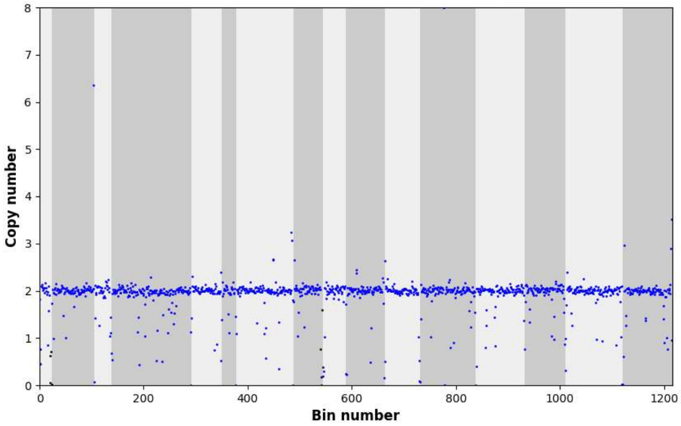


CSH908

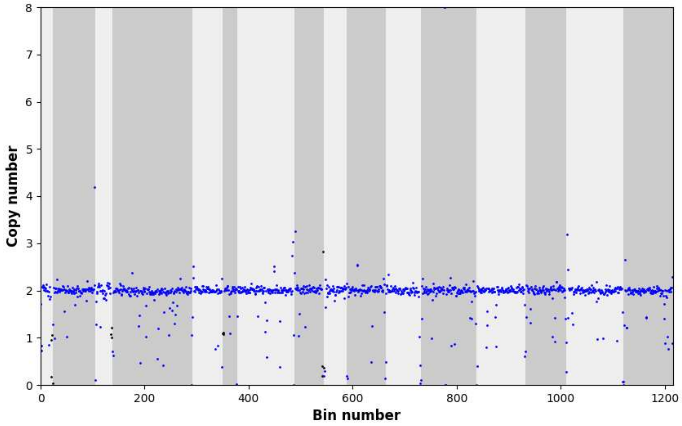


CSH909

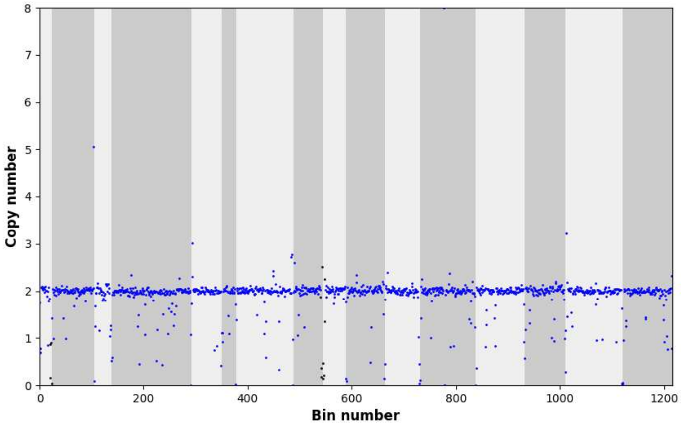


CSH910

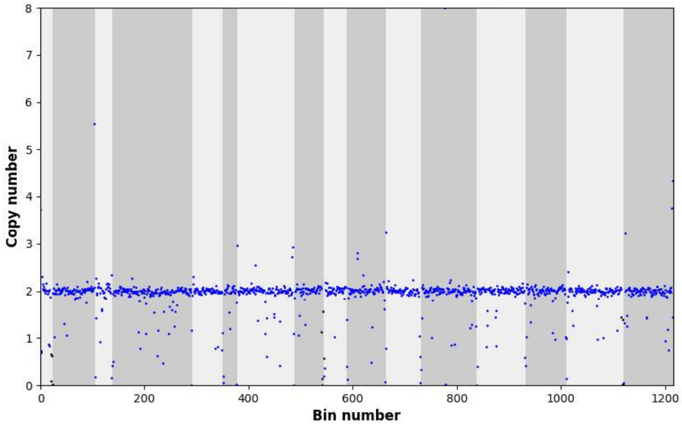


CSH911

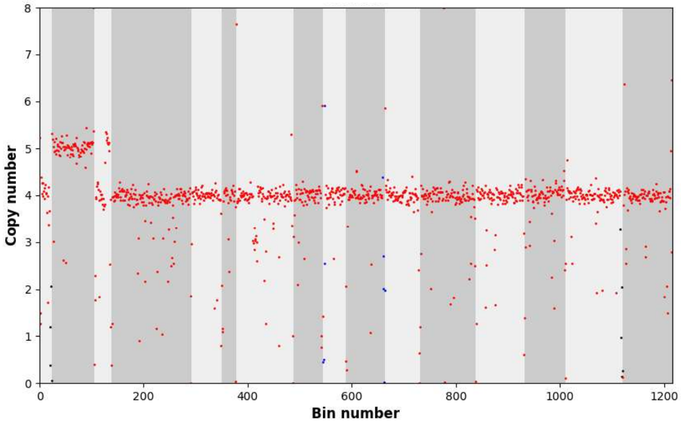


CSH913

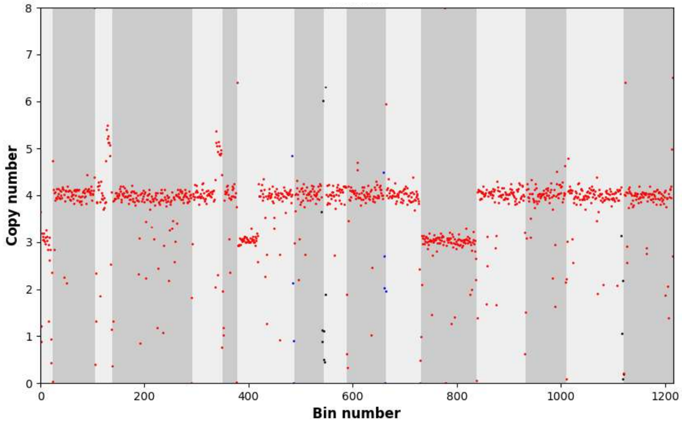

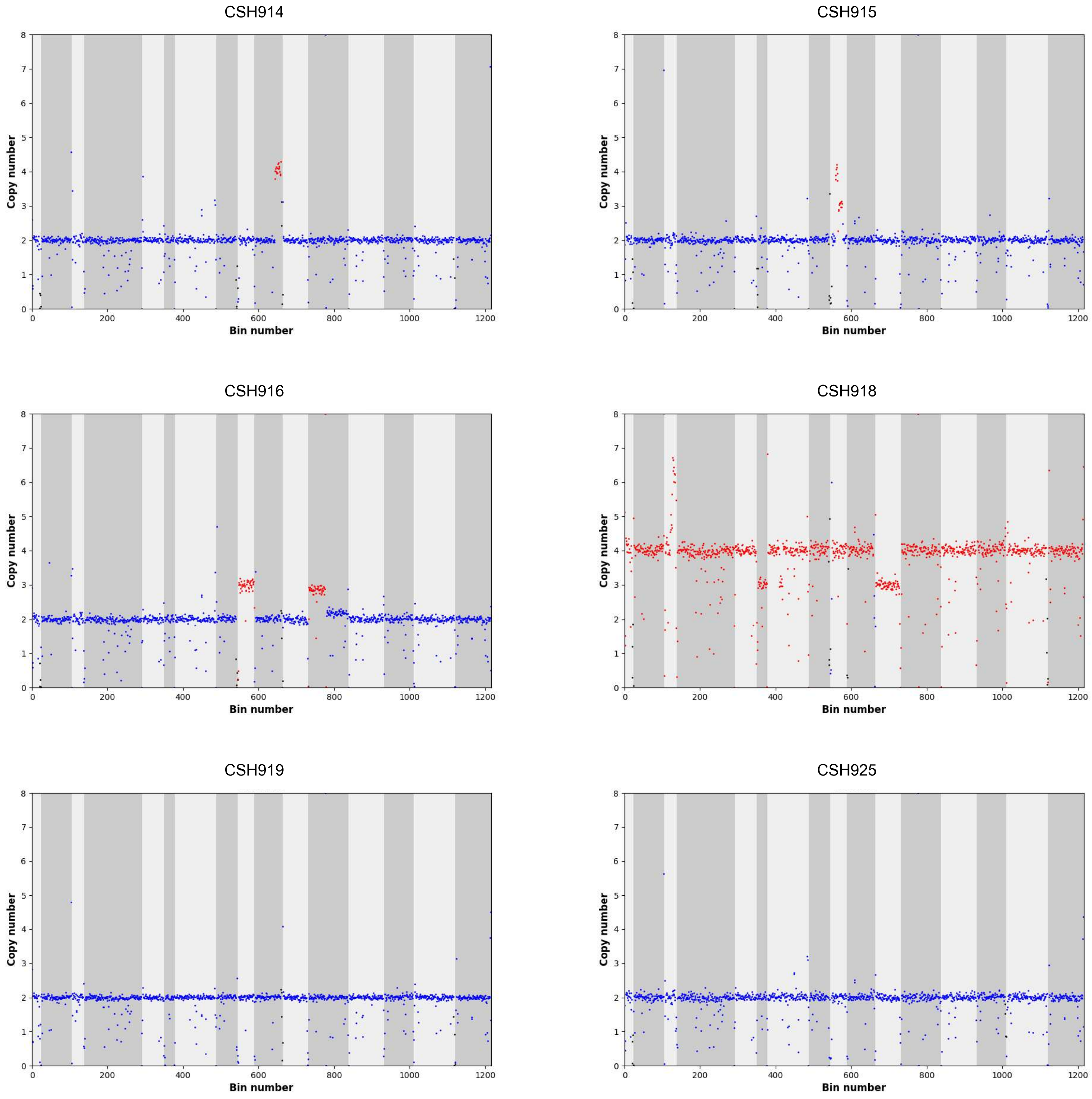
CSH914
CSH915
CSH916
CSH918
CSH919
CSH925
Copy number
Bin number
0
200
400
600
800
1000
1200
0
1
2
3
4
5
6
7
8

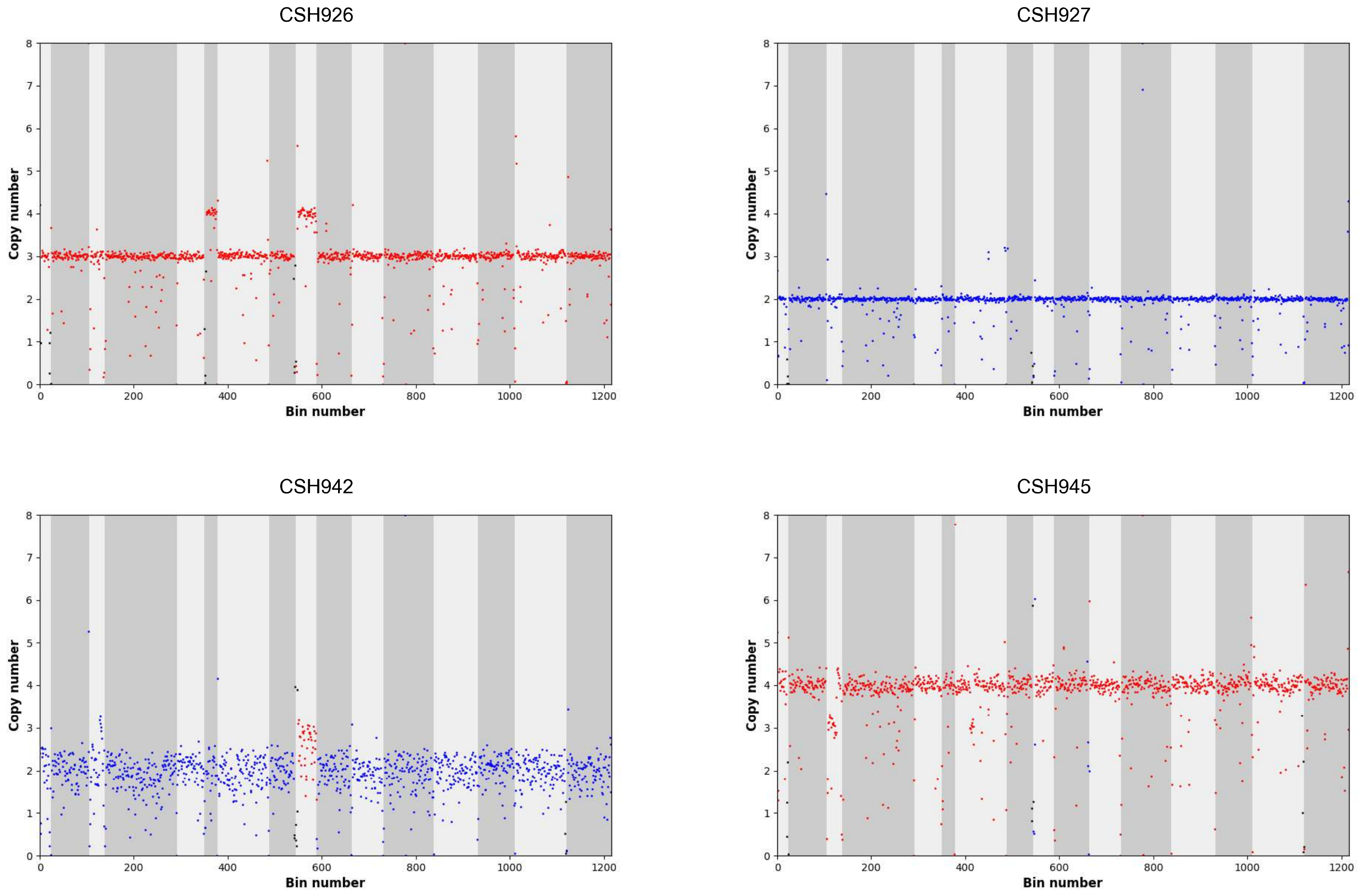


Figure S11. Ploidy and aneuploidy estimated from sequencing depth for 40 newly sequenced isolates

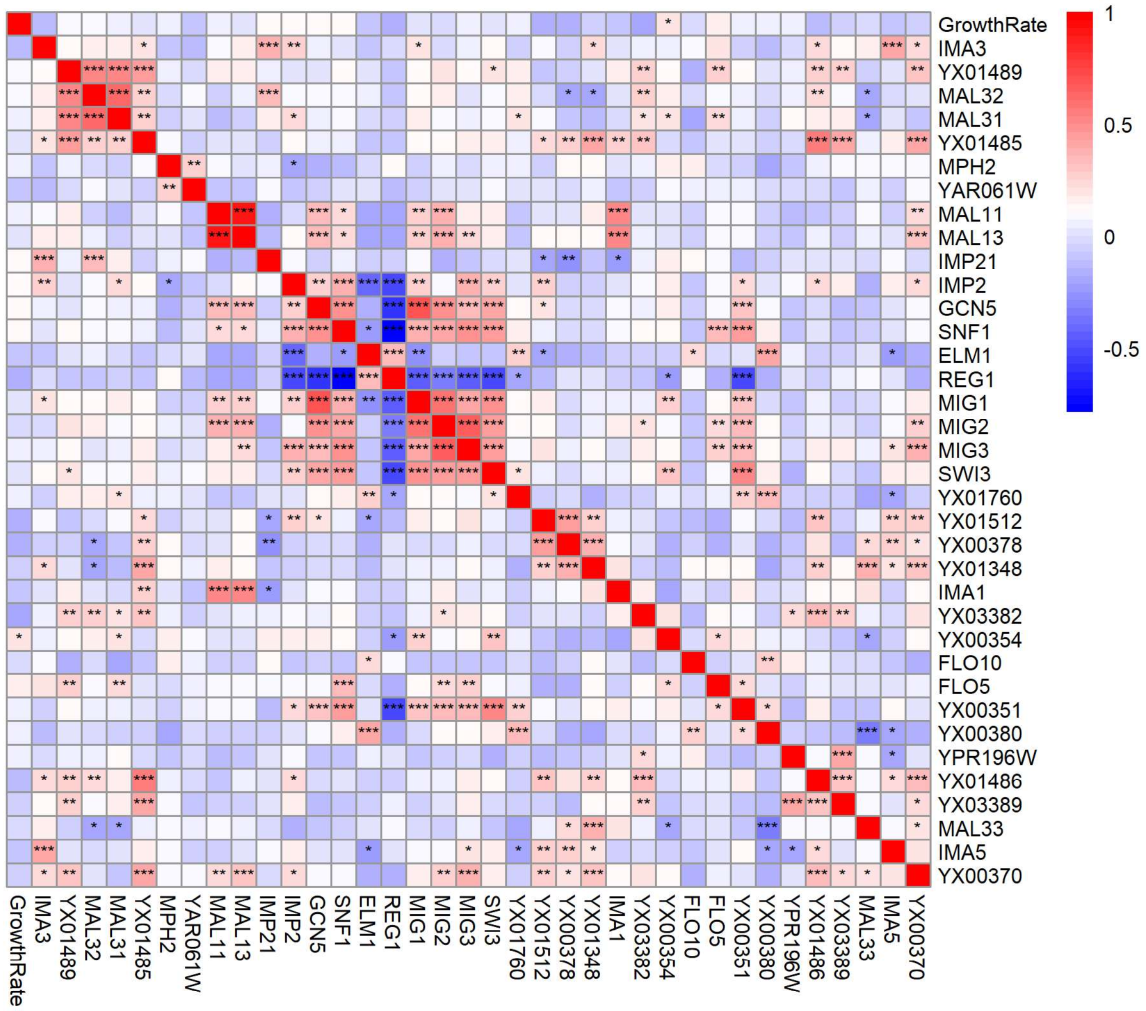


Figure S12. Spearman correlation matrix of phylogenetically independent contrasts among median growth rate in maltose media (r) and haploid normalized copy number of maltose-associated pangenome genes. Rows correspond to growth rate in maltose (GrowthRate) and gene names. Cells are colored according to strength of correlation, with positive correlations in red and negative correlations in blue. Significant correlations are denoted by asterisks. An increasing number of asterisks and color intensity represent increasing significance and strength of correlation, respectively.

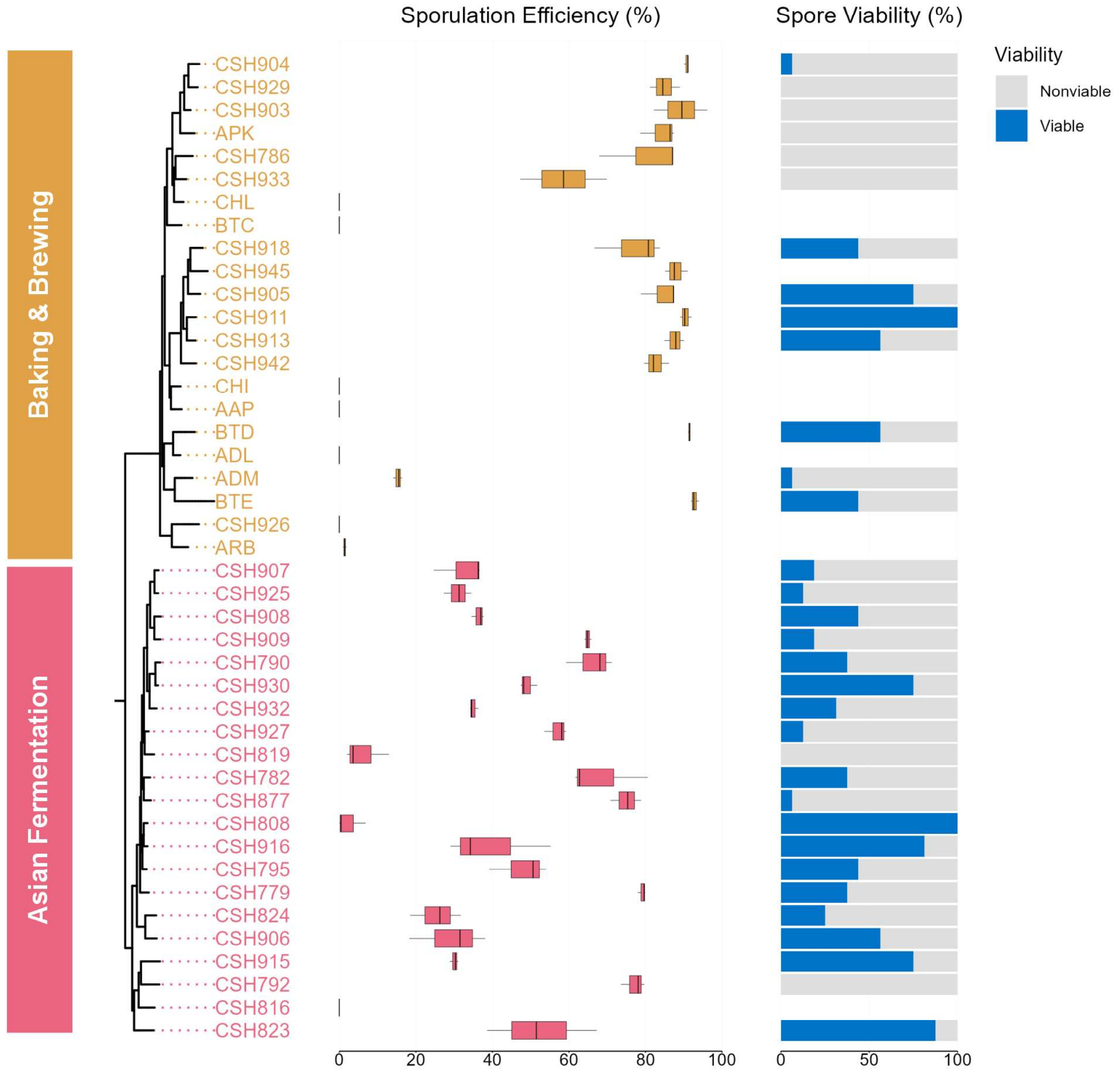


Figure S13. Sporulation efficiency and spore viability measured for 43 baking-associated isolates from Commercial Baking and Asian Fermentation groups.
